# An LFP-based characterization of population-level heterogeneity during motor preparation & execution

**DOI:** 10.64898/2026.11.09.764815

**Authors:** Michael DePass, Stephan Quessy, Ali Falaki, Numa Dancause, Ignasi Cos

**Affiliations:** Departament de Tecnologia i Comunicacions, Universitat Pompeu Fabra, Carrer Ramon Trias Fargas 25-27, 08005 Barcelona, Catalonia, Spain; Departament de Matemàtiques i Informàtica, Facultat de Matemàtiques i Informàtica, Universitat de Barcelona, Gran Via de les Corts Catalanes 585, 08007 Barcelona, Catalonia, Spain; Département de Neurosciences, Faculté de Médecine, Université de Montréal, Montréal, Québec H3C 3J7, Canada; Institut de Neurociències (UBNeuro), Universitat de Barcelona, Barcelona, Catalonia, Spain; Institut de Matemàtiques (IMUB), Universitat de Barcelona, Barcelona, Catalonia, Spain; American Physiological Society, 20852 Rockville, Maryland, USA

## Abstract

The goal of the present work is to characterize mesoscale heterogeneity of population-level neural activity in the motor areas during motor preparation and execution using high-frequency local field potentials (HF-LFPs). Understanding the distribution of neural activity at this scale is necessary since ensembles of neural populations underlie most behaviors. HF-LFPs have emerged as a promising signal for such analysis due to their stability and relationship with cortical population dynamics. Here we analyzed HF-LFPs in non-human primates performing a reach-to-grasp task. We quantified three complementary aspects of heterogeneity: variance, latent dimensionality, and directional coherence. Heterogeneity varied by region and neuromotor state, and was specific to the motor behavior being implemented, even in regions exhibiting high correlation. In addition, more heterogeneous activity tended to better encode neuromotor states. Namely, regions and neuromotor states exhibiting higher variance and dimensionality yielded superior motor decoding performance. Further analyses involving low-dimensional linear projections of the signals revealed rich intra-region interactions between ensembles of populations. The present work helps establish intra-region heterogeneity as a multi-faceted and informative measure of neural activity, even on small spatial scales. It also establishes HF-LFPs as an effective signal for capturing complex intra-region dynamics, making it a suitable candidate for multi-region, systems-level analysis.

**Significance Statement:** Though the motor system has most often been investigated through analyses of spike patterns of individual neurons, most behaviors require coordination at the level of neural populations. Recent work suggests HF-LFPs capture cortical population-level dynamics, making them apt for the characterization of neural activity at this scale. Here we analyze HF-LFP from two non-human primates, recorded during a reach-to-grasp task. Our results reveal intra-region, population-level activity that evolves over the course of motor preparation and execution. Moreover, we show that the heterogeneity of population-level activity is multifaceted and associated with motor decoding accuracy. These results are of potential interest both for future research of population-level activity and for BCI applications.

## Introduction

Traditionally, correlates of motor function have been viewed through the lens of single unit activity (SUA) (Mountcastle 1969, Georgopoulos 1982, Evarts 1983, Schieber and Hibbard 1993, Keller 2010, Lemon 2021). This work provided a detailed characterization of the somatotopic distribution of task-relevant SUA in the motor cortices. Nonetheless, implementing motor behavior extends beyond the activity of single units, often requiring the coordination of neural populations spanning multiple brain areas (Alexander 1990, Churchland and Shenoy 2024, Khilkevich 2024). Thus, understanding motor behavior requires characterizing the activity of neural populations, which according to the *population view* of the brain (Sadtler 2014, Gallego 2017, Ebitz 2022), should be considered the fundamental units of brain computation. The question of how to characterize neural correlates of behavior at this scale, however, remains.

Recent work has shown that the high frequency component of LFPs (HF-LFPs) reflects cortical, spiking population dynamics and that this activity is heterogeneous, even on small spatial scales (Gallego-Carracedo 2022, DePass 2022). Moreover, in 2019, Trautmann et al. successfully replicated SUA results in three macaque reaching studies, demonstrating that aggregate signals could represent population spiking activity with only minor distortions (Trautmann, 2019). So, while single-unit resolution is required in certain contexts, recent evidence has shown the granularity of HF-LFPs captures correlates of motor behavior while exhibiting improved temporal stability (Stark 2007, Gallego 2020). Similar signals such as multi-unit activity and spike-band power have even proven capable in complex tasks such as speech generation (Card 2026). On the opposite end of the granularity scale, scalp EEG provides a macro-scale readout of brain states which, largely, does not capture intra-region heterogeneity. This resolution has proven sufficient for applications ranging from neurological disorder diagnosis to sleep staging (Mirchi 2022). Likewise, whole brain models have successfully replicated significant aspects of spontaneous brain activity despite representing regions as homogeneous (Deco and Kringelbach 2026). While impressive, key aspects of neural dynamics (e.g., ignition-like dynamics) could only be reproduced after incorporating intra-region heterogeneity (Joglekar 2018). Similarly, capturing intra-region heterogeneity has also proven beneficial for decoding visual and motor-related brains states (Liao 2014, Kanth and Ray 2020). Thus, while capturing the vast heterogeneity at the level of single units is not required in some contexts, treating entire regions as homogeneous results in significant information loss. Together, these findings suggest HF-LFPs may serve as an apt middle ground for analyzing behavior since they capture heterogeneous, population-level activity.

While heterogeneity of neural activity at the single-unit level is well documented, the population level is underexplored (Churchland and Shenoy 2007). We know heterogeneity of SUA is beneficial for coding at the population level, but it is unclear whether the same principle applies at the mesoscale (i.e., whether more heterogeneous population activity improves multi-population coding; Chelaru 2008, Dickey 2013, Wang 2024). We hypothesize that, like SUA, population activity is heterogeneous even on small spatial scales, which enables rich coding of motor behavior within brain regions. Given this, and the fact brain regions are differentially recruited across motor behaviors, we predict the heterogeneity of population activity will be region and behavior dependent. We also predict motor behavior will be better decoded from more heterogeneous population activity. Finally, we predict that averaging activity across populations will result in information loss and, therefore, a significant reduction in motor decoding performance.

In summary, HF-LFPs capture population activity which exhibits better temporal stability than SUA while capturing informative intra-region activity, unlike EEG. We believe population-level activity, as captured by HF-LFPs, may serve as an apt middle ground for many tasks including the identification of correlates of motor function. For this reason, in the present work we characterize the heterogeneity of the population-level activity underlying motor behavior and determine it’s relationship with motor decoding.

## Methods

### Participants

The present study involved two healthy adult male rhesus macaque monkeys (Macaca mulatta), who performed a precision grip task. All experimental and surgical procedures were performed as per the guidelines stated by the Canadian Council on Animal Care and were approved by the Comité de Déontologie de l’Expérimentation sur les Animaux (CDEA) of the Université de Montréal.

### Surgical procedures

Surgical procedures were performed under sterile conditions. The monkeys were anesthetized using ketamine hydrochloride (Ketaset; 15 mg/kg; Pfizer Inc., New York, NY) and were kept anesthetized during the course of the surgery, after tracheal intubation, with approximately 2% isoflurane (Furane; Baxter, Deerfield, IL) in 100% O2. A peripheral venous catheter was used to continuously deliver Ringer’s lactate solution (10 mL/kg/h) to maintain hydration and administer any intravenous medication if needed. Before starting the surgery, a dose of buprenorphine (Temgesic; 5 µg/kg; Schering-Plough, Kenilworth, NJ) was injected subcutaneously and a dose of carprofen (4 mg/kg; Rimadyl, Zoetis Canada Inc., Kirkland, QC, Canada) was injected intramuscularly to control postsurgical pain and inflammation. Intramuscular injection of atropine (atropine sulfide; 0.04 mg/kg; Rafter 8 Products, Calgary, AB, Canada) and dexamethasone 2 (Dexacort 2; 0.5 mg/kg; Rafter 8 Products, Calgary, AB, Canada) were also given to reduce potential salivation, and brain inflammation and swelling due to craniotomy and array implantation procedures. At the beginning of the craniotomy, mannitol 20% (1,500 mg/kg; Fresenius Kabi Canada Ltd., Richmond Hill, ON, Canada) was slowly administered intravenously to further prevent brain swelling. Vital signs, including heart rate, respiratory rate, respiratory CO_2_, and arterial oxygen saturation, were monitored throughout the surgery while the body temperature was kept at 36.5C-37C using a self-regulated heating pad (Harvard Apparatus, Holliston, MA). At the end of the surgical procedure, the animal was injected with a dose of antibiotic Baytril (Enrofloxacin; 5 mg/kg; Bayer Inc., Mississauga, ON, Canada) intramuscularly to prevent infection. Additional postoperative care consisted of oral administration of Baytril and Carprofen for two days, and Gabapentin (10 mg/kg; Pfizer Inc., New York, NY) given twice per day for three days following the surgery.

To enable multi-region neural recording, craniotomies and durectomies were carried to expose the frontal cortex of both hemispheres. Five Utah microelectrode arrays (4.2 × 4.4 mm; 96 1.5 mm electrodes; Blackrock Microsystems, Salt Lake City, UT) were implanted permanently over the hand and finger representations in the left primary motor cortex (M1) and in the dorsal premotor cortex (PMd) and ventral premotor cortex (PMv) of both hemispheres (Supplemental Fig S1; all Supplemental material is available at https://doi.org/10.6084/m9.figshare.16436133). Sulcal landmarks including the arcuate and the central sulciwere used to guide the placement of electrode arrays. Short intracortical microstimulation (ICMS) trains, consisting of 13 monophasic cathodal pulses of 0.2 ms duration delivered at 350 Hz, were used to test few exploratory cortical sites to confirm the placement of arrays. ICMS trains were delivered at 1 Hz with a maximum intensity of 60 µA (Dancause et al., 2008) using a custom-made borosilicate glass-coated tungsten electrode (impedance, ∼0.5 MΩ; FHC) mounted on a micromanipulator (model 2662, David Kopf Instruments, CA, United States) to lower and insert it perpendicular to the cortex of each site. Prior to delivering the ICMS, the isoflurane was turned off to transition the animal into ketamine-diazepam sedation (Ketaset, ∼10 mg/kg/h; Valium, ∼0.01 mg.kg/h). Using these short ICMS procedures, an array was implemented in the left M1 medially to the sites that evokes orofacial responses. To implement arrays in the premotor cortex, areas medial (PMd) and lateral (PMv) to the arcuate spur were stimulated until identifying a site in each area that evokes distal movements. Arrays were implanted in PMv and PMd over these sites. Following each array implementation, wires of the implanted array were temporarily secured over the skull using bone cement (PALACOS; Heraeus Medical Components, White Bear Lake, MN).

The dura over the central sulcus of the right hemisphere was left intact and a 20 mm diameter cylindrical chamber, made out of polyether ether ketone (PEEK), was placed permanently on top of the dura over the right M1 and primary somatosensory (S1) and sealed with a removable cap. This chamber provided access to the hand representation in ipsilesional M1 for manipulations planned in the frame of a series of experiments that followed data collections used in the present study. Following array implantation, the areas of dura mater removed during durectomy were covered with Duragen (DuraGen Plus; Integra LifeSciences Holdings Corp., Princeton, NJ) and bone cement. Array connectors were housed in an acrylic box embedded within the bone cement on top of the animals’ head. Finally, anchoring surgical screws and bone cement were used to fix the entire assembly (the head post, chamber, and acrylic box) to the skull.

The dura covering the central sulcus of the right hemisphere was kept intact to implement a 20-mm in diameter cylindrical chamber made of polyether ketone over it.

### Topological references

Although the stereotaxic coordinates of the electrodes were not available, the brain areas in which each array was implanted were known. Moreover, a photograph was taken of the left hemisphere after each Utah array was implanted. This was used to design a simple planar topological map (Fig 3A), used as a referential map to draw our conclusions. Though each array contains 96 electrodes, only 32/64 from each array were used for the present analyses.

### Behavioral task

In daily experimental sessions, the monkey was brought to the laboratory to perform a reach-to-grasp task, performed while recording neural data. The monkey sat in a custom-made primate chair situated in front of the task apparatus (Fig 1A). The chair had removable panels on both sides that allowed arm movements and interactions with the task. In the task, green light emitting diodes (LEDs) on each side of the apparatus as well as inside the slots on the bottom side of the force sensors signaled the monkey which hand should be used. These LEDs were also used as GO signals, indicating to the monkey when to initiate the reach movement. The monkey then reached a target located 28 cm in front of the animal and 9 cm above the start position. In subsequent trials, the target was randomly presented in four different angles (0, 45, 90, and 135), requiring different degrees of forearm pronation and supination to orient the hand. To get the juice reward, the animal was required to reach with the instructed arm and to press force transducers in the target with the index and thumb using a precision grip. For the current analyses, only trials with 0° orientation of the grip from one session were considered. The task was operated and the data were collected with a TuckerDavis Technologies (TDT) system (Tucker-Davis Technologies Inc., Alachua, FL) using two RZ2 BioAmp processors and custom-made software. Our data acquisition equipment allowed us to simultaneously record neural activity from 256 channels from different brain areas. In addition to the data collected through the TDT system, the behavioral task was video recorded at 30 fps using a computer webcam placed on the left side of the animal. Each trial began when the animal placed both hands in the home plates (Fig 1A). Home plates contained an infrared sensor to detect the presence of the hands and signal the start of the trial. After a variable delay period between 0.1 s and 1 s, the left or right pair of LEDs was turned on (trigger Cue on) priming the monkey about the upcoming reach-to-grasp task and the instructed (active) hand to perform the task. After a variable delay (0.3–1.2 s), LEDs turned off (trigger GO), which acted as the GO cue for the monkey to initiate the task. The animal was required to initiate a self-paced reach toward the target, place his thumb and index finger in the slots and apply force (Grasp Onset) with precision grip within a maximum time window of 1.5 s. The infrared sensors embedded in the home plate detected the time point when the animal moved the hand out of the home plate, marked as the onset of reach. The onset of grasp (Grasp Onset) and the offset of grasp (Grasp Offset) were later detected by a customized MATLAB code (The MathWorks Inc., Natick, MA) as the point in time when the magnitude of force reached above and dropped below 1% of its maximal value during that trial. For this purpose, force data were low-pass filtered at 20 Hz using a zero-lag filter. To receive a juice reward, once the monkey applied forces above a prefixed threshold, he needed to keep the force above this threshold for at least 0.2 s, after which an auditory signal (beep) informed the monkey about the end of trial and receiving of juice reward. Figure 1B represents the trial progression and subdivisions (i.e., epochs) of the reach-to-grasp movement. The force threshold was set based on the force levels that the animal was comfortable to produce during training sessions. Hearing the beep, the monkey released the grip (Grasp Offset) and brought back his hands on the home plates to trigger the initiation of the next trial. The intertrial interval was between 1 s and 3 s depending on whether the orientation of force transducers was changed for the next trial. Throughout the trial, the inactive hand needed to stay rested on the home plate. To facilitate further analyses, each trial was split into five sections, called epochs, including Baseline, Pre-reach, Reach, Grasp, Post Grasp (Fig 1B) that were verified offline using webcam video recordings. During the experimental session, the NHP performed 25–40 trials with each arm and grasp orientation (a minimum of 100 trials with each arm). In the present analyses, only right-handed reach-to-grasp trials with 0° Precision grip orientation were used.

**Figure 1.**
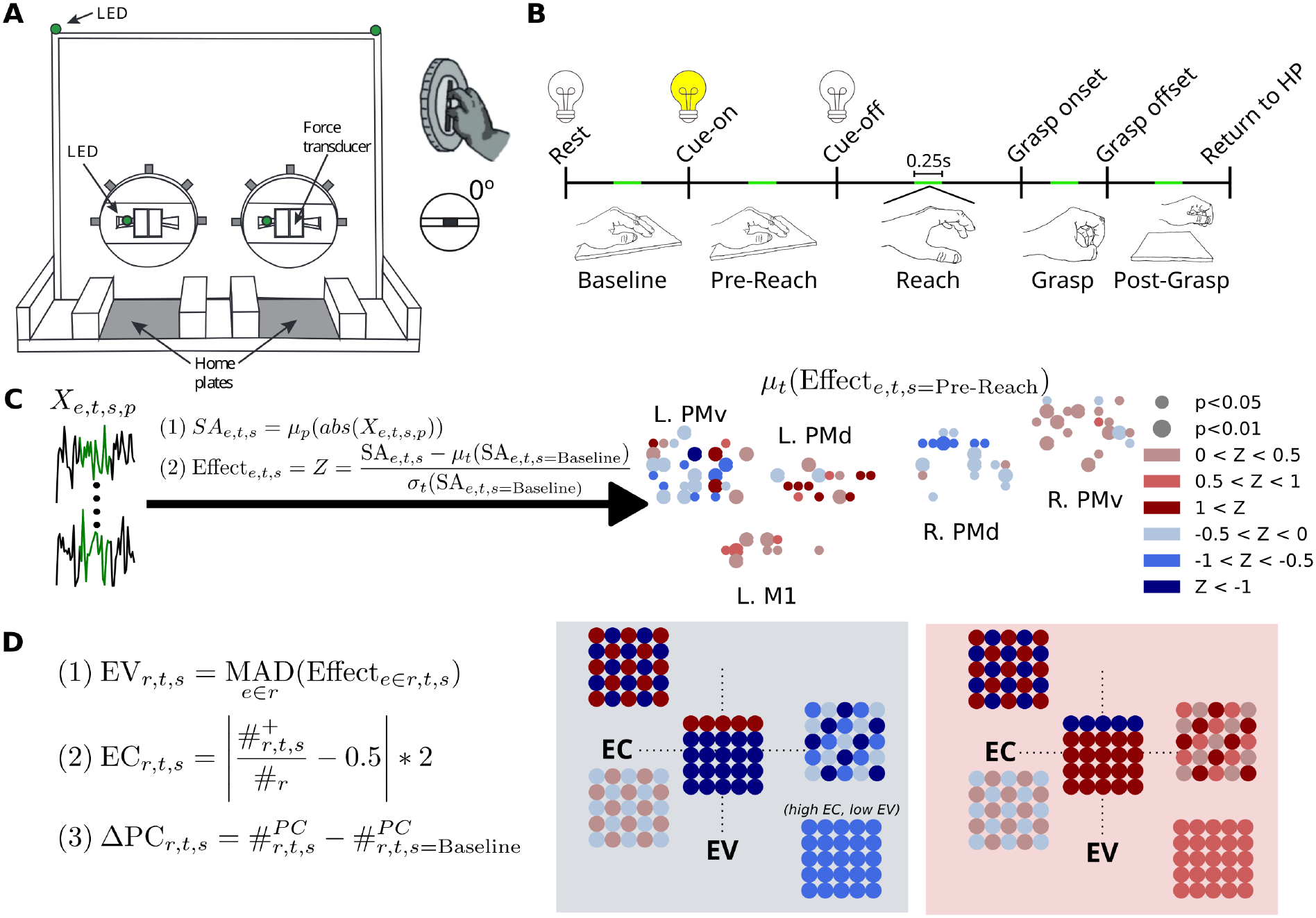
Experimental methods. **A)** Experimental apparatus used for reach-to-grasp task. Only right arm trials were analyzed. **B)** Trial timeline illustrating how the 5 neuromotor states are defined. Green segments represent the analyzed 0.25s sample of LFP taken from the midpoint of each epoch. **C)** Equations and schematic illustrating how Effect is computed from HF-LFP (Χ*_e,t,s,p_*). The green segments of the signal represent the 0.25s sample taking at the midpoint of each epoch. In the schematic, only electrodes exhibiting significant Effects (Mann Whitney U test p-value < 0.05) are shown. Circle radius indicates p-value. Circle color indicates whether the Effect was an increase (red) or decrease (blue) relative to Baseline. **D)** Heterogeneity metrics definitions (left) and illustrative examples (right). Effect Variance (EV) is the median absolute deviation (MAD) of Effect. Effect Coherence (EC) is computed from the number of electrodes exhibiting positive Effects 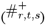 and the total number of significant Effects in the region (#_r_). Change in latent dimensionality (ΔPC) was defined as the difference in the number of principal components required to explain 66% of variance in each state relative to Baseline.

LFP data were collected from M1 and PMd of the left hemisphere, as well as PMv of both hemispheres using chronically implanted high-dimensional Utah electrode arrays (see Fig 3A for an electrode schematic). Since only right-handed movements were analyzed, we refer to regions via their laterality (e.g., cPMd instead of L. PMd). LFP data were sampled at 2,034.5 Hz and recorded digitally using our TDT system. On top of the LFP, band-pass filtered (100–500 Hz) neuronal spiking activity was recorded from the same 256 channels at 24,414 Hz. The starting time of spikes on all channels and 1,228 ms spike waveform were recorded by the data acquisition system. Spikes were detected automatically using 4 standard deviations (STDs) above the baseline noise, which was detected at the beginning of the experiment (verified by visual inspection and manually modified accordingly if needed) and locked in place throughout the duration of the experiment. For the current study, we used spiking information to de-spike the LFP data using linear interpolation. In addition to the neuronal data, applied force to each of the force transducers, sampled at 4,882.8 Hz, and various time points denoting when LEDs turned on and off, when the monkey left the home plate, and when the monkey received the juice reward and trial ends were recorded.

### Data analysis

#### Preprocessing

The first pre-processing steps were performed at the Université de Montréal with custom-written MATLAB and Brainstorm scripts (Tadel 2011). The processed data were then shared with the Math-Neuro Lab at the Department of Mathematics & Computer Science of the Universitat de Barcelona for further analysis. A second set of custom-written MATLAB scripts were used to aggregate and epoch the data. All subsequent analyses were performed with custom-made Python scripts.

First, band-pass filtering (zero-phase) between 1 and 500 Hz was done to mitigate the effects of both low and high-frequency electrical noise. Then, a notch filter was used to remove line noise, and by-connector (each connector corresponds to 32 electrodes) independent component analysis (ICA), paired with visual inspection, was performed to remove independent components corresponding to noise or physiological artifacts (e.g., head movements causing jerky movements of connectors used for neural data recording). ICA and visual inspection were repeated until no noisy ICs were visually detected. The clean ICs were then divided into segments of 4 s duration (from 2 s before grasp start to 2 s after grasp start), each representing a trial. Finally, to remove spiking activity, 3-ms windows associated with spikes were removed and replaced via linear interpolation. Trials were aligned at grasp start which was defined as the time point at which the force reached 1% of its maximal value during that trial (time zero corresponds to this grasp start time). Each trial was then divided into five behavioral epochs or neuromotor states: 1) Baseline: the period while the hand was rested in the home plate until the lights indicating trial start were illuminated; 2) Pre-reach: the period from lights on to lights off (lights off served as the GO cue indicating the animal could begin reaching for the target); 3) Reach: GO to grasp onset; 4) Grasp: 0.25 s period following grasp onset; 5) Post-grasp: the period following Grasp until the end of the trial (Fig 1B). To ensure fair sampling, samples of 0.25 s duration were extracted from each of the neuromotor states. For Grasp, this 0.25 s sample started at grasp onset. For the other states, however, samples were taken from the midpoint of the time periods described above. 40 trials from Monkey 1 and 25 trials from Monkey 2 were analyzed. The number of trials per animal differed since trials containing behavioral anomalies were removed.

#### Feature extraction

We represented the LFP dataset as a four-dimensional array *Χ* ɛ ℝ*^E×T×S×P^*, where *E* is the number of recording electrodes, *T* is the number of trials, *S* is the number of neuromotor states, and *P* is the number of sampled time points. Each entry *Χ_e,t,s,p_* corresponds to the recorded signal from electrode *e* during trial *t*, during neuromotor state *s*, at time point *p*. The SciPy Python library was then used to apply Butterworth band-pass filters to the LFP (SciPy, 2020}. LFP were analyzed in each of 8 physiologically relevant frequency bands: theta (0.5-4 Hz; abbrev. θ), alpha (8-15 Hz; abbrev. α), beta (15-30 Hz; abbrev. β), low gamma (30-70 Hz; abbrev. L. γ), high gamma (70-100 Hz; abbrev. H. γ), low ripple (100-150 Hz; abbrev. L. Rip.), high ripple (150-200 Hz; abbrev. H. Rip.), and multi-unit (200-500 Hz; abbrev. MU)). In the present work, the term “high-frequency LFP” (HF-LFP) refers to the MU band of the LFP. The first feature, Spectral Amplitude (SA), defined as the median of the rectified time series, was computed for every electrode, trial, and motor state (Fig 1C). SA served as the primary metric for subsequent classification and regression analyses. SA is similar to entire spiking activity (ESA) which boasts more robust and accurate motor decoding relative to single-unit spiking activity in certain contexts (Ahmadi 2021). SA is attractive due to its simplicity and stability over time. Moreover, SA can be computed whether or not spikes are present due to its threshold-less nature. SA is also less sensitive to high amplitude noise compared to power (power scales with the square of the signal). Zero-lag Pearson correlation (ρ) was also computed between all electrode pairs for every trial and motor state.

#### Task-relevant electrodes

Only task-relevant electrodes were analyzed. Task-relevant electrodes were determined as follows. First, SA was computed from the MU band of the LFPs. Then, binary classification between Baseline and each post-Baseline state was performed for each electrode individually (i.e., single-feature binary classification). Logistic regression classifiers, as implemented in the Scikit-learn Python library (Pedregosa 2011), were fit to the data and 50 train-test splits were performed (20% of data was left out for testing for each train-test split). Finally, permutation testing was used to determine whether the binary classifications yielded accuracy greater than chance-level where chance-level was 0.5 and α was 0.05 (Fig S1). The neuromotor state labels were shuffled 200 times for each train-test split. Electrodes were considered task-relevant if accuracy exceeded chance for any of the four binary classification tasks (one for each post-Baseline state). For Monkey 1, 250 of the 256 electrodes yielded above-chance classification accuracy in at least one of the four binary classification tasks and for Monkey 2, 246 of the 256 electrodes (Table S1).

#### Pearson correlation analyses

A single task-relevant electrode was chosen from near the center of each array/region. Pairwise Pearson correlation (ρ) was then computed between each center-point electrode and all other electrodes in the array. Mean comparisons were performed via Mann-Whitney U test as implemented in the Scipy Python library (SciPy, 2020).

#### Motor state classification

The five motor states were classified using logistic regression via the Scikit-learn Python library (Pedregosa 2011). Fifty train-test splits were performed. For each split, 80% of the data was used to train the logistic regressor and 20% was left out for testing. Train-test splits were performed such that the proportions of each class (i.e. neuromotor state) were preserved. Neuromotor states were classified using three different feature sets: 1) spectral amplitude (SA), 2) Pearson correlation (ρ), and 3) intra-region heterogeneity (i.e., EV, EC, & ΔPC). For motor state classification using heterogeneity features, heterogeneity metrics were computed from each region independently, from each region independently and then combined/concatenated (“all regions” condition – Fig 6A), and from all task-relevant electrodes across all regions (“global” condition – Fig 6A). Accuracy was recorded for every train-test split. Confusion matrices were row-normalized, yielding by-class sensitivity values on the diagonal. Sensitivity, also referred to as recall or true positive rate, is the ratio of true positives to the total number of positives in the data (e.g. the ratio of correctly classified Grasps to the total number of Grasps in the data). Statistical significance was assessed via permutation testing (see Statistical analyses).

#### Movement time regression

Movement time (MT) was defined as the interval from the GO signal to Grasp onset. Regression was used to assess the association between MT and SA. MT regression was performed for each region independently using SA from either the Pre-reach or Reach states. Moreover, forward stepwise regression was used to determine the best subset, in terms of adjusted-R^2^, of electrodes in each region for MT decoding. The best subset was the one yielding the highest adjusted-R². The regression results for the best subset of electrodes in each region are reported in Supplemental Tables S2 & S3 for Pre-reach and Reach, respectively.

#### Heterogeneity metrics

We defined three complementary metrics of intra-region heterogeneity: Effect Variance (EV), Effect Coherence (EC), and latent dimensionality relative to Baseline (ΔPC). The first two metrics, EV and EC, are both computed from the Effect which we defined as the Z-scored difference in SA for each post-Baseline state relative to Baseline (Fig 1C). Effect was calculated on a per-trial basis for each electrode. e, t, s, and P refer to electrode number, trial number, neuromotor state, and time point, respectively. Effect Variance (EV) was then derived from Effect*_r,t,s_* using the median absolute deviation (MAD) to reduce the influence of outliers, noisy trials, etc. EV was calculated on a by-region basis, thus `r` refers to the set of electrodes in a particular region. Finally, Effect Coherence (EC) was also derived from Effect. EC is proportional to the ratio of the number of electrodes in a region exhibiting increases in SA relative to Baseline, 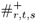, to the total number of electrodes in the region, #_r_ (Fig 1D). If we denote this ratio as R, then EC=2|*R*−0.5|. Thus, an EC of 1 means all significant changes (i.e., Effects) in SA are in the same direction (either all positive or all negative) and an EC of 0 means there is an even split of positive and negative changes. EC for a particular trial, brain region, and neuromotor state, was only calculated if at least 3 electrodes exhibited Effects that were at least 0.25 standard deviations above (or below) the Baseline mean. These checks ensured that EC could not be dominated by one or two electrodes and that electrodes were only included if their corresponding SA values differed markedly from Baseline.

The third metric of heterogeneity, ΔPC, was the difference in latent dimensionality of the HF-LFP time series (i.e. number of principal components required to exceed 66% of explained variance) relative to Baseline. The 66% explained variance threshold was chosen because a larger threshold (e.g., 95%) would retain many low-variance components that would likely be dominated by noise or artifact rather than representing coordinated, population-level activity. To assess threshold-dependent stability of the ΔPC metric, we repeated the analysis with the 0.60, 0.70, and 0.75 thresholds (Table S4). Unlike EV and EC, ΔPC is calculated directly from the HF-LFP time series rather than from the Effect. ΔPC was also computed per trial for all neuromotor states and brain regions. In summary, we intended these 3 metrics to provide complementary descriptions of intra-region heterogeneity: EV is a measure of spread, EC is a measure of directional agreement (i.e. coherence) of the sign of the Effects within a particular region, and ΔPC is a measure of the latent dimensionality of the LFPs relative to Baseline. Higher EV and ΔPC imply higher heterogeneity while high EC implies lower heterogeneity.

#### Granger causality analysis

Principal component analysis (PCA) was performed on the cleaned LFPs for each combination of brain region and motor state. PCA yielded a dimensionality reduced (with respect to the number of electrodes) projection of the original neural data. Enough principal components (PCs) were kept to explain at least 66% variance. Granger causality (GC) was then computed via the grangercausalitytests function from the Statsmodels Python library (Seabold 2010). A max lag of 2 time points (∼1 ms) was chosen to capture short-latency directed activity while avoiding overfitting. Granger causality was then computed for every trial between all pairs of sources. We adopted a conservative, consensus-based approach to identify reliable, directed interactions. GC between sources was considered significant only if it reached p < 0.05 (χ2 test) in at least 75% of trials. In this way, we prioritized replicability across trials though weaker or less stable GC connections may be omitted.

#### N features analysis

Feature importance was assessed using the LGBMClassifier function from the LightGBM Python library (Shi 2025). Features were considered more important the more they appeared across all trees in the fit model. Once feature importance was established, motor states were classified (using the LGBMClassifier) for an increasing number of features (from 1 to 10), in order of importance. Three conditions were considered for this analysis: “all features”, “all pc”, and “one pc per region”. The “all features” condition refers to the use of the original feature set (i.e., electrode space). The “all pc” condition, however, involves the use of the original features projected onto the first `n` principal components (as determined by PCA). Enough components were kept from each brain region to explain 66% variance. Finally, the “one pc per region” condition involved keeping a single feature/source from each brain region. The above analysis was performed for both the SA and the ρ feature sets.

## Statistical analyses

Regression analyses were performed using the Statsmodels Python library (Seabold 2010). Depending on the data structure in each analysis, we employed either ordinary least squares models (OLS) with cluster-robust standard errors or linear mixed-effects models. For OLS regression models, we used t-tests to assess the significance of model coefficients. For linear mixed-effects models, we used Wald z-tests to evaluate model coefficient significance.

For the Pearson correlation model, we clustered within electrode pairs to account for the dependence between repeated measures across frequency bands. For the Effect, EV, EC, and ΔPC regression models, we clustered standard errors by trial to account for dependencies within each reach-to-grasp movement sequence. For the sensitivity vs. heterogeneity analysis, we used a linear mixed-effects model. Since we obtained 4 sensitivity values (one per post-Baseline motor state) from the same classifier, within-region observations were dependent. To account for this, we included regions as random intercepts in the model. Regarding trial aggregation prior to regression, our approach differed between the correlation and heterogeneity analyses. We aggregated trials via averaging prior to correlation regression to reduce the complexity of an otherwise massive model. For heterogeneity regression, however, we kept trial-level measurements of EV, EC, and ΔPC since we considered them to be biological replicates.

In addition to regression models, nonparametric pairwise comparisons were performed via Mann-Whitney U test. Bonferroni corrections for multiple comparisons were also applied (reported p-values are adjusted unless otherwise stated). We used the iPMv (i.e., R. PMv) brain region as a reference to define the region categorical variable. The iPMv was selected since it was the only ipsilateral region present in both animals.

We used permutation tests to assess the statistical significance of classification accuracy (i.e. motor decoding performance). For each train-test split, the data labels in the training set were shuffled 500 times. This generated a null distribution of classification accuracy for each train-test split. The p-value for each split was then considered to be the percentage of accuracies from the null distribution that exceeded the accuracy of the classifier fit to the unshuffled data. The median p-value across splits was then calculated and compared to an ɑ of 0.05 to determine the overall statistical significance of the classification. Permutation tests were one-sided.

The same permutation testing method described above was also used to report by-state statistical significance in the confusion matrices. Only statistical significance of the diagonal of the confusion matrices is reported.

## Results

We studied intra-region heterogeneity of population-level neural activity in the motor areas during motor preparation and execution in a reach-to-grasp task performed by two macaque monkeys. The monkeys were implanted with microelectrode arrays in the ventral and dorsal, contralateral and ipsilateral premotor areas (cPMv/iPMv and cPMd/iPMd, respectively) and the contralateral primary motor area (cM1). The recording setup allowed for the analysis of signals in a band ranging from 1 to 500 Hz. Each trial was split into five epochs, from rest and motor preparation to execution, which we refer to as neuromotor states. The data allowed us to investigate the heterogeneity within each brain region for every neuromotor state. We quantified heterogeneity via three complementary metrics: EC, EV, and ΔPC (see Methods). We then used these metrics to provide a multifaceted characterization of intra-region heterogeneity of brain coding during the motor task.

### Pair-wise correlation decreases with frequency in the brain areas with best decoding performance

We were interested in identifying relationships between the intra-region correlation profiles and motor decoding performance (i.e., neuromotor state classification accuracy). Motor decoding performance was only calculated for the MU frequency band because it yielded the best classification accuracy in previous work and due to its high correlation with population activity (Stark 2007, Gallego-Carracedo 2022, DePass 2022).

We could not establish an overall relationship between absolute intra-region correlation and decoding performance (Fig 2). For Monkey 1, for example, MU correlation was lowest in the cPMd and highest in the cM1 (Fig 2A, left panel); however, LFPs from both areas yielded excellent motor decoding performance (Fig 2B, left panel). Another example is the similar correlation exhibited by cPMd and iPMv in Monkey 1 (Mann-Whitney U test, U=891, p_uncorrected_=0.942; Fig 2A, left panel), and their stark difference in decoding performance (accuracy_cPMd_=0.95, accuracy_iPMv_=0.302). iPMv even fails to exceed chance level (permutation test, p=0.09). Similarly, iPMv MU correlation and cM1 MU correlation did not differ significantly despite their stark difference in motor decoding accuracy (Mann-Whitney U test, U=1023, p_uncorrected_=0.711_;_ accuracy_iPMv_=0.302 for iPMv and accuracy= 0.731 for cM1). For Monkey 2, while the specific correlation profiles differed somewhat from Monkey 1, there was still no clear relationship between correlation and decoding accuracy. Moreover, while Monkey 2 LFPs were generally less correlated, decoding accuracy did not differ markedly between animals.

**Figure 2.**
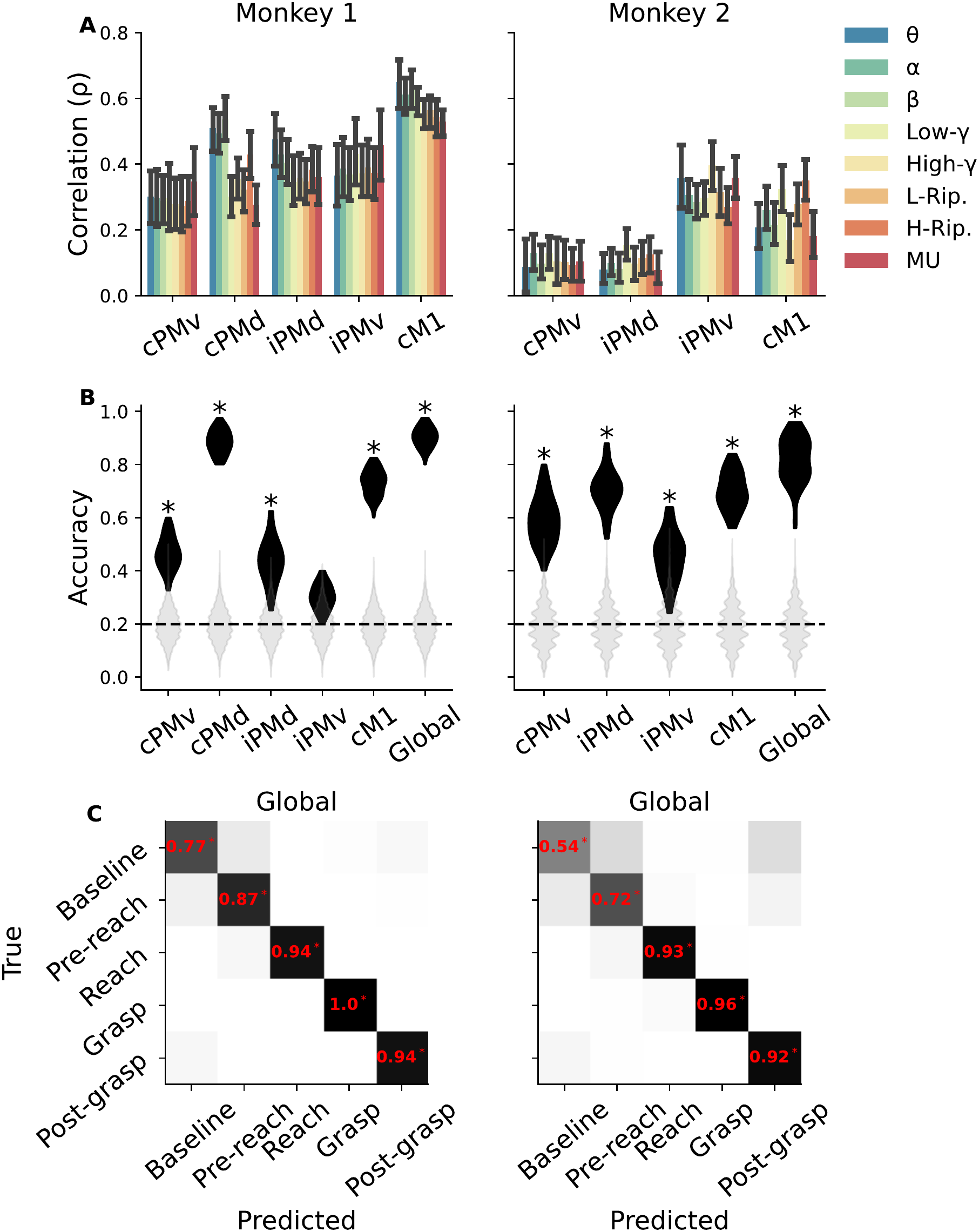
HF-LFP is less correlated than low frequency LFP in regions that yielded the best motor decoding performance. **A)** LFP intra-region correlation bar plots for eight frequency bands for Monkey 1 (left) and Monkey 2 (right). **B)** Classification accuracy violin plots for all brain regions for Monkey 1 (left) and Monkey 2 (right). Asterisks indicate statistical significance relative to chance-level (dashed line). Grey violins represent the null distribution of classifier accuracy as determined by permutation testing. **C)** Global (i.e. all areas), row-normalized confusion matrices for Monkey 1 (left) and Monkey 2 (right). By-class sensitivity is shown in red on the diagonal and statistical markers indicate significance as determined by permutation testing. Significance is only reported on the diagonal. Darker cells indicate a larger number of classifications in the “True label” - “Predicted label” pair.

Regions that yielded the best motor decoding performance tended to exhibit significantly lower correlation in the higher frequency bands. In Monkey 1 cPMd and cM1, for example, correlations in the MU frequency band were lower than in all other bands (Fig 2A, left panel). The correlation in the MU band was roughly 0.2 and 0.1 lower than in the θ band for cPMd and cM1 (Mann-Whitney U test, U=116 & U=222, p=0.005 & p=0.22, for cPMd & cM1, respectively; p_uncorrected_=0.0003 for cM1). It is worth nothing, however, that low within-region correlation did not guarantee good decoding performance. The correlation within the MU band in Monkey 1 cM1 (Fig 2A, left panel) was the highest of any region, despite yielding a better decoding performance than cPMv, iPMd, and iPMv.

Our decoding analyses confirm that HF-LFPs (200-500 Hz) contain plenty of information about motor preparation and execution. Specifically, HF-LFPs from all regions yielded above chance-level classification accuracy except for Monkey 1 iPMv (Fig 2B). Moreover, the confusion matrices in Fig. 2C show that all five neuromotor states could be identified with near perfect accuracy using the global (i.e., from all regions) signal. Likewise, all five states could be accurately identified using signals from single brain areas, from Monkey 1 cPMd and Monkey 2 iPMd (Fig S2).

Though decoding performance was generally well above chance, it varied significantly between animals. This may not be surprising, given that arrays only provide signals from a relatively small sample within each recorded region. The iPMd provides the clearest example of this with Monkey 1 iPMd yielding an accuracy of 0.44 while Monkey 2 iPMd boasted an accuracy of 0.70 (Mann-Whitney U test, U=23, p<0.0001). These findings suggest population-level neural activity in the motor cortices encodes states pertaining to motor preparation and execution in a widely distributed, bilateral, region-specific, and frequency-specific manner. Moreover, the fact that certain regions exhibited markedly different levels of motor decoding performance between animals might also suggest a high level of intra-region heterogeneity. If population activity is highly heterogeneous within region, one would expect even small differences in array placement or subject-specific anatomico-functional distribution to exert a significant impact on decoding performance.

### HF-LFPs were associated with movement time

Though neuromotor states can be accurately decoded from HF-LFP (Fig 2C), it is important to determine whether the HF-LFP signals predict motor behavior. To this end, we performed regression analysis between Movement Time (MT), defined as the time interval between the GO signal and the grasp onset, and the HF-LFP time series from several cortical areas during Pre-reach and Reach. MT in both animals averaged roughly 0.5s, ranging between 0.25s and 1s.

Our results show that MT could be reliably predicted from HF-LFP-derived SA during both Pre-reach (prior to movement onset) and Reach, from several cortical regions (Table S2). For Monkey 1, MT could be decoded from cPMd (F-test, F=3.48, p=0.005, R^2^=0.364) and iPMd (F-test, F=3.335, p=0.021, R^2^=0.193). The SA from three electrodes in the cPMd and three electrodes in the iPMd was associated with MT. Interestingly, both regions contained a mix of electrodes that were either positively or negatively associated with MT. For Monkey 2, both cPMv (F-test, F=5.05, p=0.016, R^2^=0.252) and iPMd (F-test, F=3.814, p=0.039, R^2^=0.19) SAs were significant predictors of MT. A single electrode in each region was associated with MT. Note that MT was not predicted by iPMv signals from either animal.

Overall, this analysis shows that HF-LFPs (and, therefore, population-level neural activity) encode specific movement parameters. Moreover, the fact there was a mix of positive and negative associations between SA and MT further suggests a high level of heterogeneity across the population-level neural activity of the PMd. We repeated the analysis with HF-LFPs during Reach and found SA during Reach was also associated with MT (Table S3). One significant result that differed from the previous, Pre-reach based analysis was that SA in the cM1 was associated with MT in Monkey 2.

### HF-LFPs capture rich population dynamics on small spatial scales

To further explore intra-region heterogeneity, we then calculated the *latent dimensionality* of neural activity for each neuromotor state via PCA and determined if there were any Granger causal connections between the principal components (Fig 3). The number of principal components needed to explain most of the variance in the neural data (#PC) is thought to represent the number of patterns of coordinated firing activity, shaped by the local neural circuitry, among subsets of neurons in a population (Gallego 2017, Perich 2025, Sadtler 2014, Gallego-Carracedo 2022). Principal component analysis (PCA) in Monkey 1 revealed that latent dimensionality (#PC required to explain 66% of variance) of the HF-LFPs tended to increase relative to Baseline during both motor preparation (Pre-reach) and execution (Reach, Grasp, & Post-grasp) in the cPMv, cPMd, iPMd, and cM1 (Fig 3B, left panel). The only region which did not exhibit an increase in latent dimensionality relative to Baseline was the iPMv. These findings mirror the decoding performance within each region. The iPMv was the only region in Monkey 1 which did not yield above chance-level motor decoding performance (Fig 2B). It is worth noting that latent dimensionality did not appear to covary with the number of electrodes recorded from in the region. For example, the data consisted of 32 channels from the cPMd and cM1 for Monkey 1, while 64 channels were recorded from the cPMv, iPMd, and iPMv (Fig 3A). Despite recording from fewer electrodes in the cPMd and cM1, these regions exhibited the highest #PC of all 5 regions. It is worth noting our #PC results are in line with previous research which demonstrated latent dimensionality in PMd was 50% greater than in M1 during reaching (Gallego 2020).

**Figure 3.**
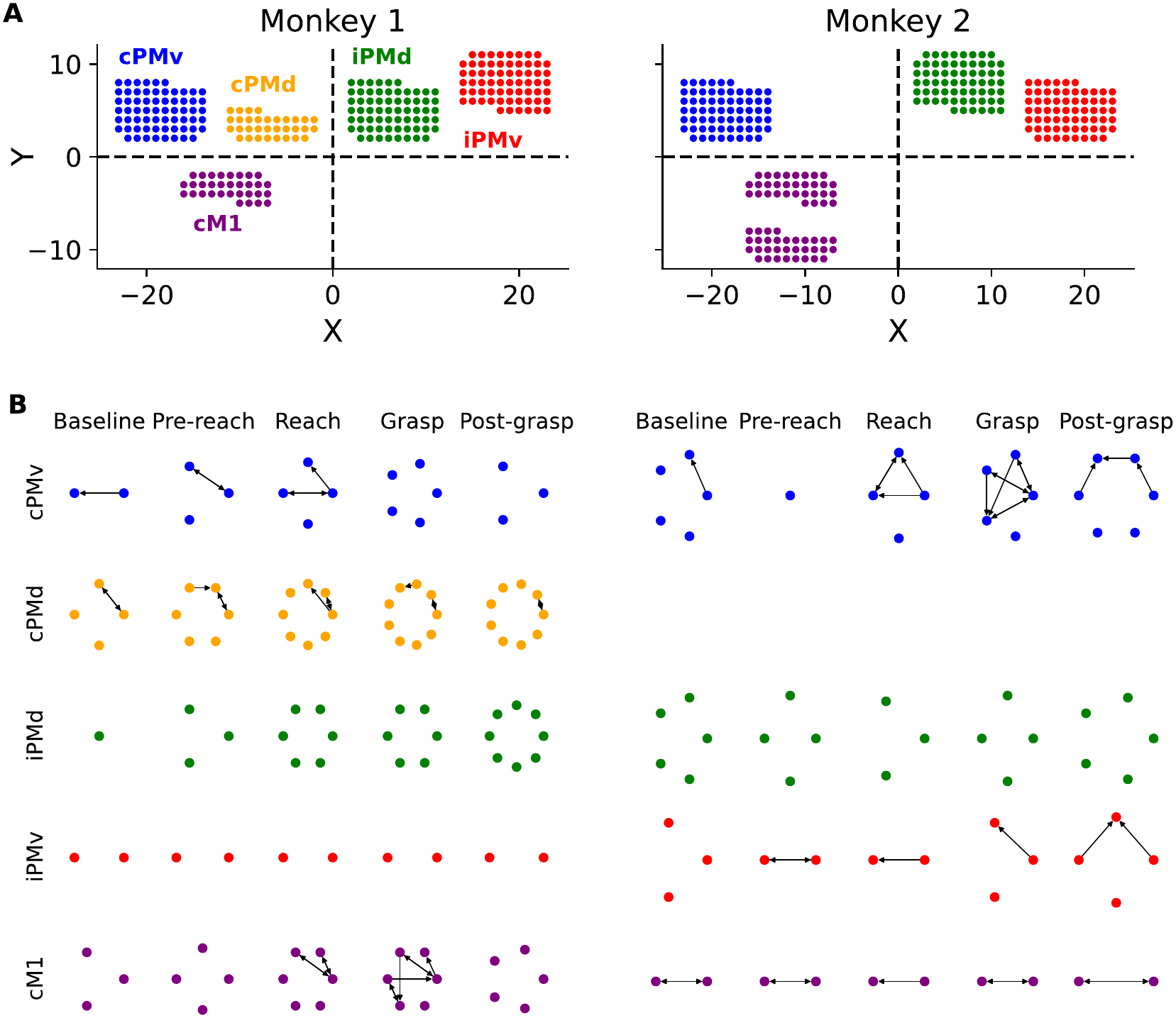
Source space Granger causality analysis reveals state and region-dependent dimensionality and connectivity. **A)** Schematics of electrode coordinates for Monkey 1 (left) and Monkey 2 (right). Relative electrode positions are representative within-array though inter-array distances, however, are not representative. Position units (i.e., X and Y) are arbitrary. The true distance between neighboring electrodes is 0.4 mm. **B)** PCA was performed on HF-LFP for each neuromotor state and brain region. The number of nodes in each graph equals the median number of principal components required to explain 66% of the variance. Edges of the graph represent significant Granger causal connections.

We then calculated Granger causality for connections between principal components within each region. For Monkey 1, Granger causal connections were only present in the regions contralateral to the moving hand (all movements were performed with the right arm/hand). Granger causal connections in Monkey 2, however, were also significant in iPMv (Fig 2B, right panel). For Monkey 2, Baseline tended to have a relatively large #PC compared to other neuromotor states with the exception of the cM1. #PCs in this area did not change throughout the entire reach-to-grasp movement. Though it is difficult to generalize from results of a small number of animals exhibiting unique profiles, the results still point to a high degree of heterogeneity of population-level neural activity in the motor areas during reach-to-grasp. Both #PC and the number of Granger causal connections were region- and state-dependent for both animals. Furthermore, these results illustrate the complex interactions taking place on small spatial scales within brain regions/arrays during the reach-to-grasp movement.

### Intra-region heterogeneity of HF-LFPs is region and state specific

To provide a more complete characterization of intra-region heterogeneity, we calculated two additional metrics complementary to the #PC relative to Baseline metric (ΔPC). These two additional metrics were derived from the Effect metric, which we defined as the mean difference in SA relative to the Baseline state (ΔSA; Fig 1C; see Methods). The Effect metric differs across neuromotor states and brain regions. In Monkey 1 (F test, F=102.5, p<0.0001, R^2^=0.466; Table 1), the Effect was positive in cPMd, indicating an increase in SA relative to Baseline during Pre-reach and Reach. However, it was negative during Grasp and Post-grasp (Fig 4A, left panel). In the case of the cPMv, however, there were no significant differences across neuro-motor states. Effect varied heavily across brain regions as well. For Monkey 2, the Effect was primarily positive in the cM1, and negative in iPMd and iPMv.

**Figure 4.**
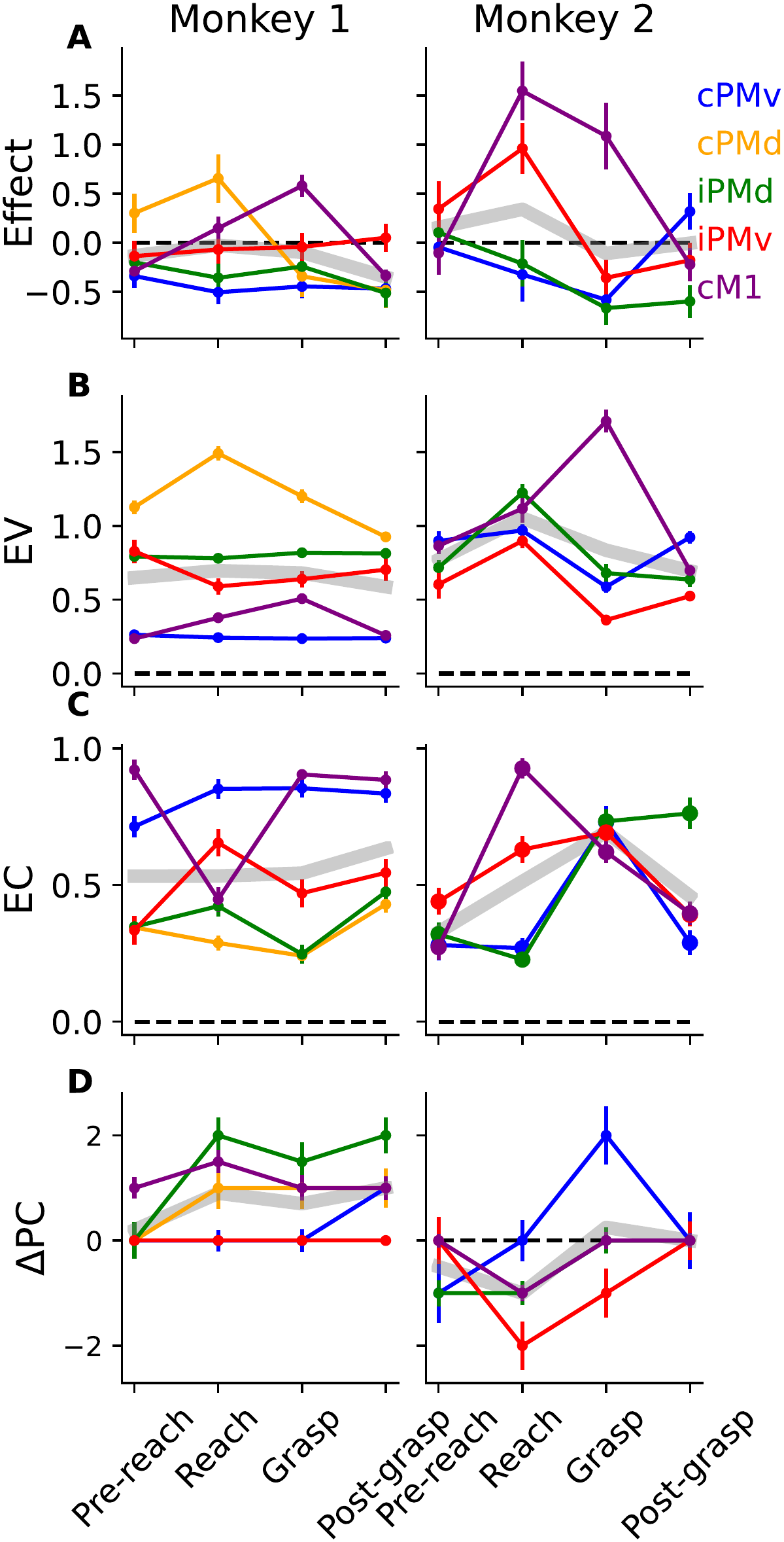
Intra-region heterogeneity (effect variance — EV, and effect coherence — EC) and latent dimensionality of HF-LFP depend on brain region and neuromotor state. **A)** Time course of the Effect (i.e. spectral amplitude relative to Baseline), **B)** EV, **C)** EC, and **D)** ΔPC. All metrics are relative to baseline. The dashed lines show Baseline and the grey lines the mean across regions. Error bars indicate the standard error of the mean.

**Table 1.** Effect (ΔSA) and intra-region heterogeneity regression results. Multiple linear regression coefficients for models predicting intra-region heterogeneity from motor state and brain region. Numbers in parentheses indicate standard error of the mean.

|  | *Effect* |  | *EV* |  | *EC* |  | *ΔPC* |  |
|---|---|---|---|---|---|---|---|---|
|  | Monkey 1 | Monkey 2 | Monkey 1 | Monkey 2 | Monkey 1 | Monkey 2 | Monkey 1 | Monkey 2 |
| Motor state: Reach | 0.00<br>(0.14) | 0.71<sup>*</sup><br>(0.31) | -0.26<sup>*</sup><br>(0.11) | 0.16<br>(0.13) | 0.18<sup>*</sup><br>(0.07) | 0.09<br>(0.08) | 0.05<br>(0.04) | -0.80<br>(0.64) |
| Motor state: Grasp | 0.08<br>(0.12) | -0.79<sup>**</sup><br>(0.26) | -0.23<sup>*</sup><br>(0.10) | -0.28<sup>*</sup><br>(0.11) | 0.10<br>(0.08) | 0.16<sup>*</sup><br>(0.07) | 0.05<br>(0.04) | 0.92<br>(0.58) |
| Motor state: Post-grasp | 0.14<br>(0.12) | -0.81<sup>**</sup><br>(0.25) | -0.16<br>(0.12) | -0.21<sup>*</sup><br>(0.10) | 0.09<br>(0.08) | -0.05<br>(0.07) | -0.00<br>(0.05) | 2.00<sup>**</sup><br>(0.61) |
| Region: cPMv | -0.19<sup>*</sup><br>(0.08) | -0.70<sup>**</sup><br>(0.22) | -0.68<sup>***</sup><br>(0.08) | 0.18<sup>*</sup><br>(0.08) | 0.19<sup>**</sup><br>(0.06) | -0.12<br>(0.08) | 0.17<br>(0.21) | 0.32<br>(0.72) |
| Region: cPMd | 0.44<sup>***</sup><br>(0.08) |  | 0.16<sup>*</sup><br>(0.08) |  | -0.09<br>(0.07) |  | 0.25<br>(0.35) |  |
| Region: iPMd | -0.08<br>(0.10) | -0.49<sup>*</sup><br>(0.24) | -0.12<br>(0.09) | 0.11<br>(0.10) | -0.08<br>(0.06) | -0.14<sup>*</sup><br>(0.06) | 0.22<br>(0.36) | 1.08<sup>*</sup><br>(0.51) |
| Region: cM1 | -0.09<br>(0.10) | -0.71<sup>**</sup><br>(0.24) | -0.72<sup>***</sup><br>(0.08) | 0.23<br>(0.13) | 0.39<sup>***</sup><br>(0.07) | -0.19<sup>**</sup><br>(0.07) | 0.50<sup>*</sup><br>(0.21) | 1.36<sup>**</sup><br>(0.52) |
| Reach x cPMv | -0.18<br>(0.14) | -0.99<sup>***</sup><br>(0.29) | 0.23<sup>*</sup><br>(0.11) | -0.05<br>(0.11) | -0.00<br>(0.08) | -0.17<br>(0.12) | 0.57<sup>*</sup><br>(0.25) | 1.40<br>(0.95) |
| Reach x cPMd | 0.33<sup>*</sup><br>(0.15) |  | 0.72<sup>***</sup><br>(0.12) |  | -0.18<sup>*</sup><br>(0.09) |  | 1.35<sup>***</sup><br>(0.32) |  |
| Reach x iPMd | -0.14<br>(0.16) | -1.08<sup>***</sup><br>(0.30) | 0.21<br>(0.11) | 0.14<br>(0.14) | -0.12<br>(0.08) | -0.23<sup>**</sup><br>(0.09) | 1.58<sup>***</sup><br>(0.36) | 0.32<br>(0.61) |
| Reach x cM1 | 0.38<sup>**</sup><br>(0.14) | 1.07<sup>**</sup><br>(0.41) | 0.41<sup>***</sup><br>(0.11) | 0.16<br>(0.19) | -0.50<sup>***</sup><br>(0.10) | 0.47<sup>***</sup><br>(0.10) | 1.10<sup>***</sup><br>(0.22) | 0.56<br>(0.70) |
| Grasp x cPMv | -0.22<br>(0.13) | 0.31<br>(0.25) | 0.19<sup>*</sup><br>(0.10) | -0.02<br>(0.09) | 0.08<br>(0.09) | 0.10<br>(0.12) | 0.68<sup>**</sup><br>(0.22) | 1.36<br>(0.92) |
| Grasp x cPMd | -0.71<sup>***</sup><br>(0.14) |  | 0.32<sup>**</sup><br>(0.10) |  | -0.20<sup>*</sup><br>(0.09) |  | 1.65<sup>***</sup><br>(0.29) |  |
| Grasp x iPMd | -0.12<br>(0.14) | 0.01<br>(0.27) | 0.25<sup>*</sup><br>(0.11) | 0.22<br>(0.13) | -0.16<br>(0.10) | 0.20<sup>*</sup><br>(0.10) | 1.25<sup>**</sup><br>(0.39) | -0.64<br>(0.72) |
| Grasp x cM1 | 0.72<sup>***</sup><br>(0.15) | 1.98<sup>***</sup><br>(0.28) | 0.50<sup>***</sup><br>(0.10) | 1.09<sup>***</sup><br>(0.17) | -0.06<br>(0.10) | 0.12<br>(0.09) | 0.88<sup>***</sup><br>(0.22) | -0.28<br>(0.61) |
| Post-grasp x cPMv | -0.29<sup>*</sup><br>(0.13) | 1.21<sup>***</sup><br>(0.26) | 0.13<br>(0.12) | 0.28<sup>**</sup><br>(0.10) | 0.06<br>(0.08) | -0.01<br>(0.09) | 0.80<sup>**</sup><br>(0.25) | -0.48<br>(1.02) |
| Post-grasp x cPMd | -0.89<sup>***</sup><br>(0.11) |  | -0.02<br>(0.14) |  | -0.01<br>(0.11) |  | 2.00<sup>***</sup><br>(0.29) |  |
| Post-grasp x iPMd | -0.43<sup>**</sup><br>(0.14) | 0.11<br>(0.27) | 0.18<br>(0.14) | 0.10<br>(0.11) | 0.07<br>(0.08) | 0.32<sup>***</sup><br>(0.09) | 2.18<sup>***</sup><br>(0.36) | -1.36<sup>*</sup><br>(0.68) |
| Post-grasp x cM1 | -0.27<sup>*</sup><br>(0.13) | 0.71<sup>*</sup><br>(0.28) | 0.18<br>(0.12) | -0.01<br>(0.13) | -0.08<br>(0.09) | 0.14<br>(0.10) | 0.48<sup>*</sup><br>(0.21) | -1.56<sup>*</sup><br>(0.68) |
| Intercept | -0.12<br>(0.07) | 0.62<sup>**</sup><br>(0.22) | 0.96<sup>***</sup><br>(0.08) | 0.72<sup>***</sup><br>(0.10) | 0.42<sup>***</sup><br>(0.05) | 0.49<sup>***</sup><br>(0.05) | -0.00<br>(0.05) | -1.60<sup>***</sup><br>(0.46) |
| Observations | 800 | 400 | 800 | 400 | 800 | 400 | 800 | 400 |
| R<sup>2</sup> | 0.48 | 0.62 | 0.69 | 0.52 | 0.40 | 0.37 | 0.18 | 0.17 |
| Adjusted R<sup>2</sup> | 0.47 | 0.61 | 0.69 | 0.50 | 0.39 | 0.34 | 0.16 | 0.14 |
| Residual Std. Error | 0.35 | 0.54 | 0.25 | 0.30 | 0.24 | 0.24 | 1.73 | 1.87 |
| F Statistic | 102.51<sup>***</sup> | 128.77<sup>***</sup> | 393.34<sup>***</sup> | 59.06<sup>***</sup> | 127.88<sup>***</sup> | 36.88<sup>***</sup> | 15.57<sup>***</sup> | 8.34<sup>***</sup> |

Note: <sup>\*</sup>p<0.05; <sup>\*\*</sup>p<0.01; <sup>\*\*\*</sup>p<0.001

The first new metric we introduce is Effect Variance (EV; a measure of spread of the neural response; see Fig 1C & Methods) was also specific to each neuromotor state and brain region (Fig 4B). EV in the cPMd exhibited much larger amplitude than in all other regions in Monkey 1 (Mann-Whitney U tests, U>92, p<0.0001^1^). In addition, EV was significantly larger in cPMd during Reach compared to other neuromotor states (Mann-Whitney U tests, U>305, p<0.0004). cPMv, however, exhibited a very low EV, which did not vary by state for Monkey 1 (Fig 4B, left panel).

The second new metric is Effect Coherence (EC; a measure of directional uniformity of the neural response; see Fig 1C & Methods), which varied with neuromotor state and brain region (Fig 4C). EC was higher in the cM1 and cPMv compared to other brain regions for Monkey 1 (Mann-Whitney U tests, U>2917, p<0.0001). Interestingly, EC in cM1 during Reach decreased sharply for Monkey 1 while it increased sharply for Monkey 2 (Fig 4C).

Relative latent dimensionality (ΔPC; a measure of the increase ^1^ in latent dimensionality relative to Baseline; see Fig 1C & Methods) also depended on neuromotor state and brain region. For Monkey 1, all regions increased their latent dimensionality (i.e. positive ΔPC) relative to Baseline except for the iPMv (Fig 4D, left panel). Moreover, increases in latent dimensionality were the most pronounced in the Reach, Grasp, and Post-grasp states. The Pre-reach state, however, did not exhibit any increase in latent dimensionality relative to Baseline. For Monkey 2, unlike for Monkey 1, regions often exhibited decreases in latent dimensionality (i.e. negative ΔPC; Fig 4D, right panel).

Overall, Effect, EV, EC, and ΔPC depended on neuromotor state and brain region for both animals. The nature of the specific dependencies between the heterogeneity metrics, neuromotor state, and brain region, however, differed between animals.

To ensure the ΔPC results were not overly sensitive to the explained variance threshold of 0.66, we computed ΔPC for thresholds of 0.60, 0.70, and 0.75 for Monkey 1 (Table S4). We found most of the results to be robust albeit with a few exceptions. ΔPC during Reach and Grasp in the cPMv, for example, increased from 0 to 1 across the thresholds. The same is true for ΔPC in the iPMd during Pre-Reach. Despite these minor differences, we believe our conclusion that ΔPC depends on neuromotor state and brain region holds.

### Spread and latent dimensionality covary with motor decoding accuracy

We first calculated EV, EC, and ΔPC metrics for each neuromotor state and brain region. Second, we assessed the association between decoding performance and our three metrics by means of linear regression (Fig 5). Sensitivity was used as the metric of decoding performance (see Methods; represented by the diagonal of the row-normalized, by-region confusion matrices – Fig S3). Regression analyses revealed a statistically significant relationship between ΔPC and sensitivity (i.e. motor decoding performance) in both monkeys (Wald tests, z=2.723 & z=4.408, p=0.006 & p<0.0001 for Monkey 1 and 2, respectively; Table 2). For Monkey 2, EV was also a significant predictor of sensitivity (Wald test, z=3.404, p=0.0006).

**Figure 5.**
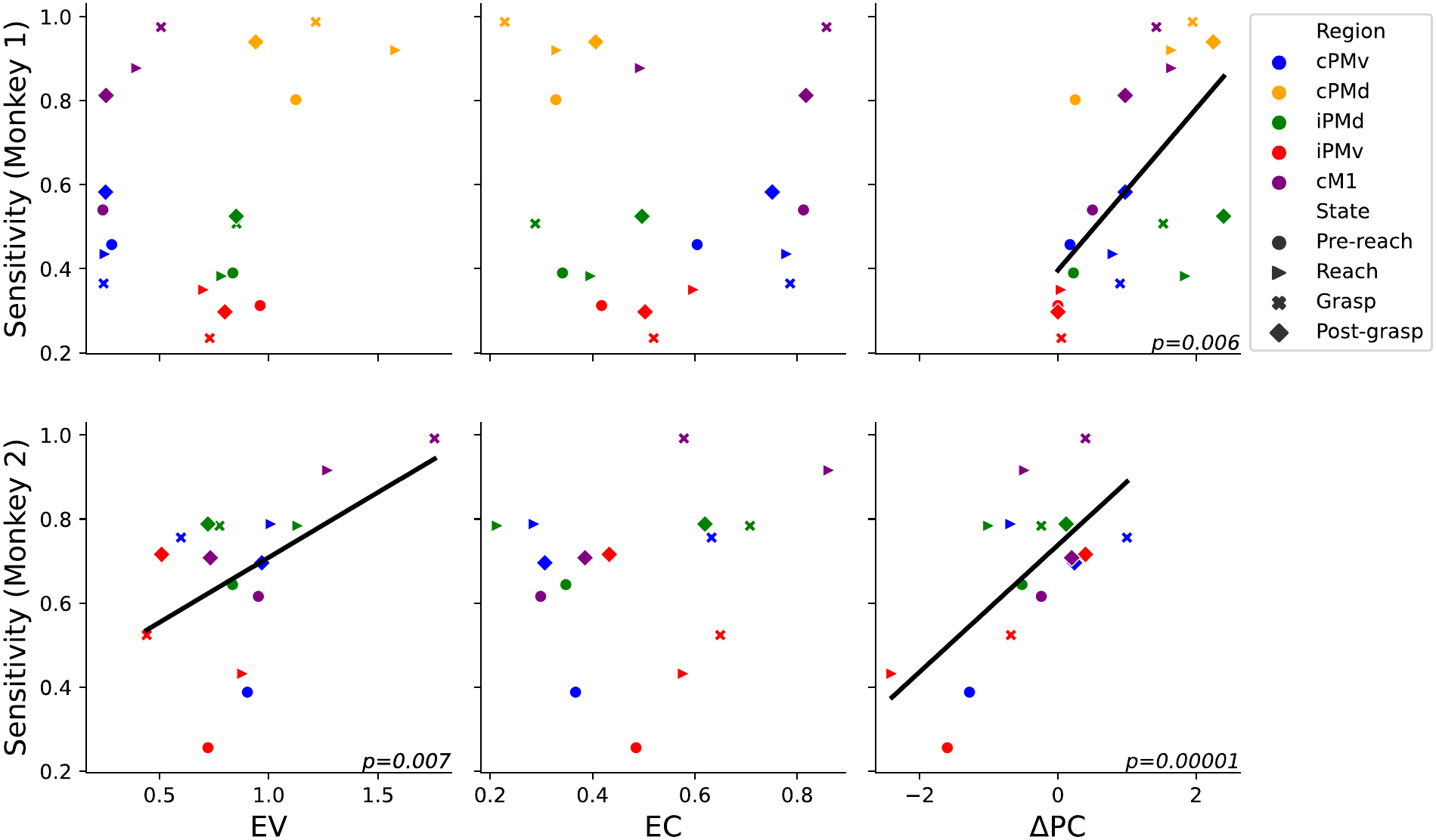
Motor decoding performance covaried with intra-region heterogeneity. Sensitivity as a function of intra-region heterogeneity (EV, EC, & ΔPC) of HF-LFP for Monkey 1 (top row) and Monkey 2 (bottom row). Sensitivity values indicate the true positive rate for each motor state (e.g., percentage of grasps accurately classified as grasps). Sensitivity values were determined by 5-state classification via logistic regression. Colors represent brain areas and shapes represent neuromotor states. Chance-level is 0.2.

**Table 2.** Regression coefficients from models predicting sensitivity from three metrics of intra-region heterogeneity: Effect Coherence (EC), Effect Variance (EV), and latent dimensionality (ΔPC). Sensitivity is a measure of classification (i.e. decoding) performance which indicates the proportion of true positives to actual positives.

|  | *Sensitivity* |  |
|---|---|---|
|  | Monkey 1 | Monkey 2 |
| Group Var | 6.16 | 0.09 |
|  | (5.30) | (0.32) |
| Intercept | 0.41 | 0.37<sup>**</sup> |
|  | (0.22) | (0.12) |
| EC | -0.07 | 0.17 |
|  | (0.23) | (0.15) |
| EV | 0.16 | 0.32<sup>***</sup> |
|  | (0.16) | (0.09) |
| ΔPC | 0.10<sup>**</sup> | 0.15<sup>***</sup> |
|  | (0.04) | (0.03) |
| Observations | 20 | 16 |
| Residual Std. Error | 0.08 | 0.10 |

Note: <sup>\*</sup>p<0.05; <sup>\*\*</sup>p<0.01; <sup>\*\*\*</sup>p<0.001

While our results suggest a positive association between EV and ΔPC and motor decoding performance, our results do not suggest EC is a reliable predictor. In Monkey 1, for example, decoding performance was highest in the cPMd and cM1 despite the regions’ stark difference in EC (Figure 5 top middle panel).

The strong statistical evidence of the dependence of decoding performance on EV in Monkey 2 and ΔPC in both monkeys suggested that neuromotor states could be identified based on the heterogeneity metrics. To validate this, we performed a classification of neuromotor states using the three metrics. The results indicate that our metrics alone were sufficient to decode neuromotor states at well above chance-level (Figure 6).

**Figure 6.**
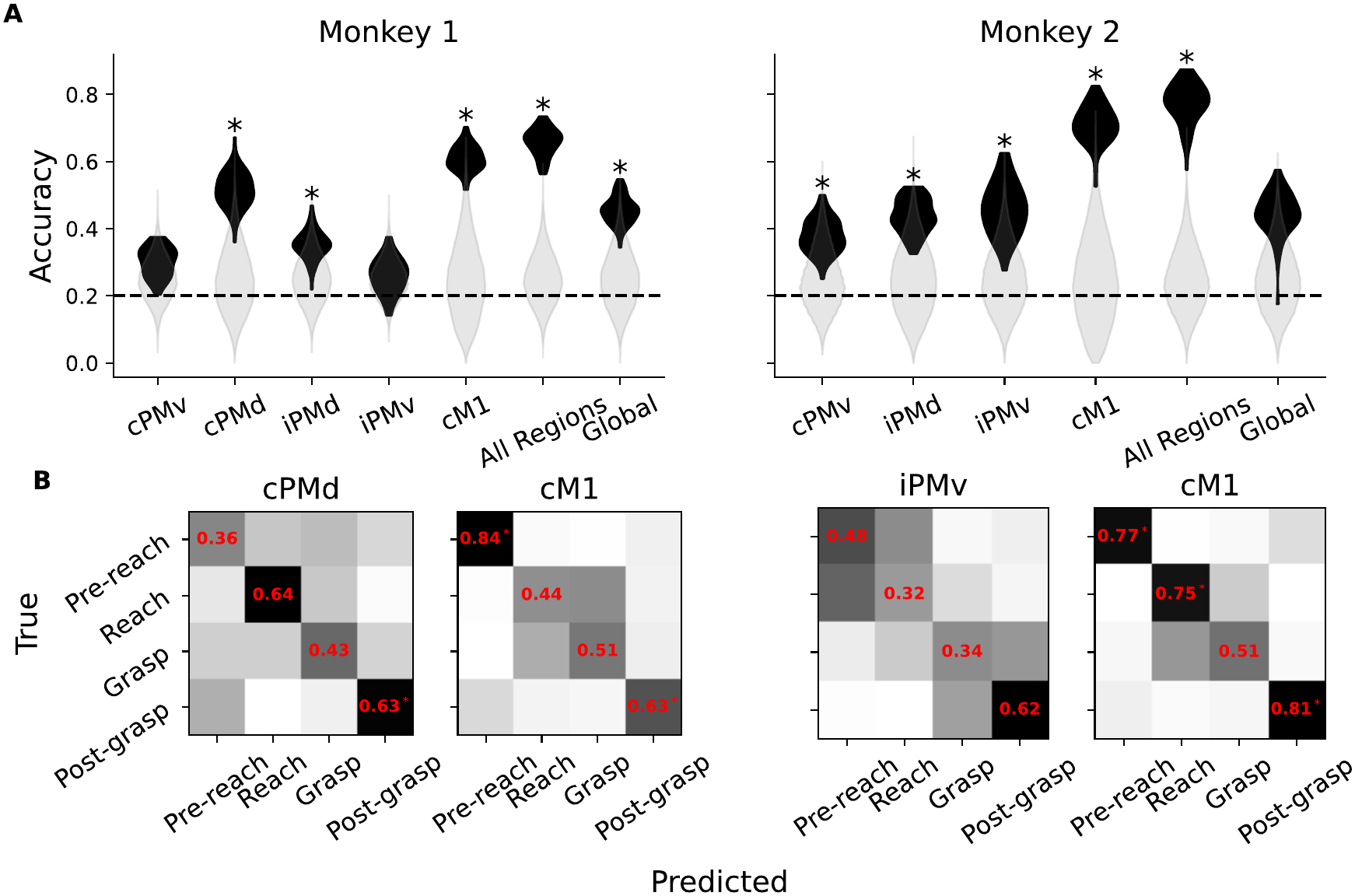
Neuromotor states could be classified from intra-region heterogeneity alone (i.e. EV, EC, & ΔPC). **A)** Classification accuracy as a function of brain area for Monkey 1 (left) and Monkey 2 (right). The “all regions” conditions involved classifying neuromotor states combining intra-region heterogeneity features from each region The “global” condition involved computing the three heterogeneity metrics on the whole dataset rather than within each region. The horizontal dashed line indicates chance-level. **B)** Confusion matrices for the two regions yielding the highest classification accuracy for Monkey 1 (left) and Monkey 2 (right). Confusion matrices are row-normalized yielding sensitivity values on the diagonal.

To reinforce the notion that intra-region heterogeneity was region and state-specific, we performed Global and All-Regions analyses (“Global” and “All-Regions” x-axis labels in Fig 6AB, respectively). Note that the “Global” case refers to analyses in which we classified neuromotor states using heterogeneity metrics computed after pooling electrodes from all regions together. The “All-Regions” case refers to an analysis in which states were classified using the concatenation of intra-region heterogeneity metrics from all regions. Our findings indicate the 5 neuromotor states could be identified from intra-region heterogeneity at well above chance-level in most brain regions. For Monkey 1, the cPMd and cM1 yielded a well above-chance accuracy (accuracy_cPMd_=0.52 and accuracy_cM1_=0.62; chance-level=0.2; permutation tests, p=0.0299 & p=0.001; Fig 6AB). The cPMd and cM1 intra-region heterogeneities yielded higher classification accuracies than the Global case (Mann-Whitney U tests, U=2266 & U=2480, p<0.0001 & p<0.0001, for cPMd & cM1 compared to Global). For Monkey 2, all conditions yielded above chance-level decoding performance except for the Global one (permutation tests, p<0.04; Fig 6A). Confusion matrices revealed that certain regions were better suited for decoding particular neuromotor states, suggesting a specific anatomico-functional distribution, consistent with several previous studies (He 1993, Galea 1994, Fluet 2010, Griffin 2015). cPMd for Monkey 1, for instance, yielded above chance-level sensitivity for Reach and Post-grasp (sensitivity_Reach_=0.65 and sensitivity_Post-grasp_=0.64; permutation tests, p_Reach_=0.035 & p_Post-grasp_=0.035). In the cM1, however, Pre-reach decoding far exceeded chance-level while it failed to exceed chance in the cPMd (sensitivity_cM1_=0.86 and sensitivity_cPMd_=0.37; permutations tests, p_cM1_=0.005 & p_cPMd_=0.16). cM1 in Monkey 2 yielded results similar to Monkey 1 in that Pre-reach and Post-grasp could be decoded above chance-level. Unlike in Monkey 1, however, Reach decoding was also significant (sensitivity_Pre-reach, Reach, Post-grasp_=0.78, 0.72, 0.84; permutation tests, p_Pre-reach, Reach, Post-grasp_=0.0225, 0.04, 0.005). Together, our results indicate that small scale, intra-region heterogeneity of population-level neural activity is a significant predictor of decoding performance. These results also indicate a multifaceted characterization of heterogeneity is beneficial since some metrics are associated with decoding performance while others are not. We repeated this analysis with a stricter channel rejection protocol (see Methods) and found the results were robust (Table S5).

### Classification accuracy plateaus after ∼5 features and PC-space features yielded reduced neuromotor state classification accuracy

To assess whether population activity as captured by HF-LFP lies at a suitable level of granularity for characterizing motor behavior, we performed neuromotor state classification with the SA and Pearson correlation (ρ) feature sets.

We used PCA to assess classification performance at various levels of granularity for both feature sets. Figure 7 shows the relationship between three levels of granularity: fine granularity (“all features”), medium granularity (“all pc”), coarse granularity (“one pc per region”) and their resulting accuracy. The “all PC” feature set consists of all PCs required to explain 95% variance. The coarse granularity case (“one PC per region”) feature set consists of a single PC for each region. For both metrics (i.e. SA and ρ) and animals, reducing the feature set to a single PC per region resulted in a significant reduction in classification accuracy (Mann-Whitney U test, U>315, p<0.0001; Fig 7). Interestingly, classification accuracy generally did not improve with the addition of features. In some cases, decoding performance barely exceeded chance-level (Figure 7 - bottom panels). Compared to fine granularity, medium granularity also resulted in decreased classification accuracy, though the magnitude of this decrease varied greatly between animals. For Monkey 1, there was a small but significant decrease in classification accuracy after projecting to the PC-space (Mann-Whitney U test, U=354006, p<0.0001; Fig 7 - upper-left panel). For Monkey 2, projection to the PC-space reduced classification accuracy much more (though it remained significantly greater than the single PC per region condition). For the ρ features, projection to the PC-space also led to a significant decrease in decoding performance though the disparity was larger in Monkey 2 compared to Monkey 1 (Mann-Whitney U tests, U>8155, p<0.0001 in all cases; Fig 7 - bottom panels). These results suggest projecting HF-LFP onto the PC-space significantly reduces decoding performance, even when controlling for the number of features. This suggests either that the highest variance PCs are not the most relevant for motor decoding, or that intra-region population-level neural dynamics are highly non-linear and, therefore, not adequately captured by a linear technique such as PCA.

**Figure 7.**
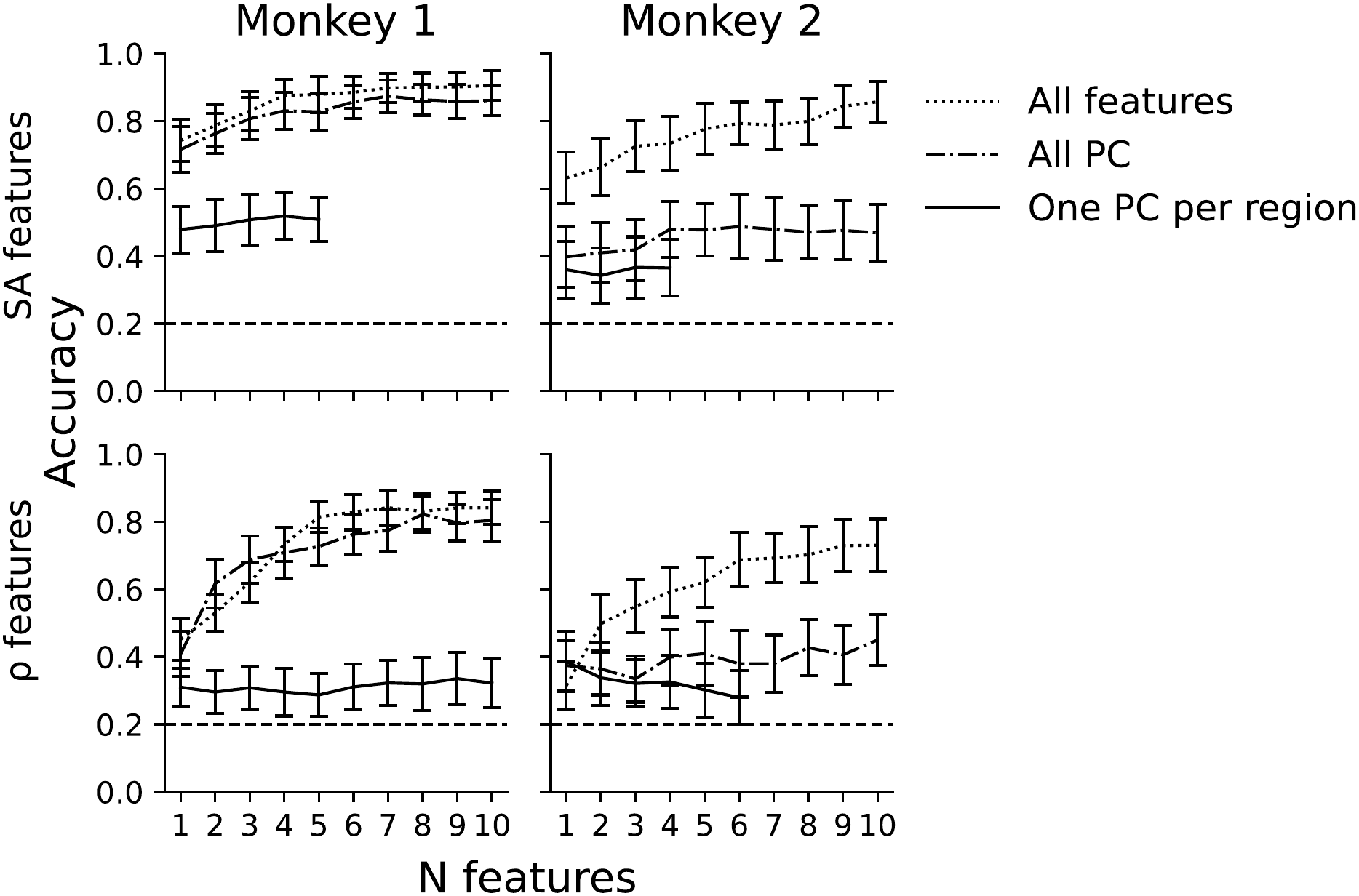
Neuromotor state classification accuracy depends on the number, kind (i.e., spectral amplitude or Pearson correlation), and degree of dimensionality reduction of the features. The “all features” condition represents classification without dimensionality reduction. The “all pc” condition involved performing classification on principal components (abbrev. PC) derived from each brain area. The “one pc per region” condition involved classification where only one PC per region could be used. The horizontal dotted line indicates chance-level.

## Discussion

We used a dataset of LFPs, recorded from the pre- and primary motor cortices of two macaque monkeys during movement preparation and execution, to investigate the heterogeneity of the underlying population-level neural activity. Our results show that correlation among LFPs within brain area decreases with LFP frequency, particularly in the areas yielding the highest motor decoding accuracies (cPMd, iPMd, and cM1). This finding, combined with the fact HF-LFPs have been shown to correlate with population spiking activity, motivated further analysis of the MU frequency band. To this end, we showed HF-LFPs capture rich intra-region interactions across ensembles of neural populations throughout the reach-to-grasp movement (Fig 3). We then employed three complementary metrics to quantitatively assess heterogeneity: spread (EV), directional agreement (EC), and latent dimensionality (ΔPC). Regression analyses revealed brain-region and neuromotor-state specific profiles of population heterogeneity. We then performed an additional analysis to determine whether motor states were better encoded in more heterogeneous population activity. We found that, while EC did not exhibit any significant relationship, EV and ΔPC were positively correlated with motor decoding performance. We then sought to determine whether intra-region heterogeneity was more informative than global heterogeneity. Classification analyses revealed that intra-region heterogeneity is, in fact, more informative in terms of motor decoding performance. Finally, to further assess whether the mesoscale resolution provided by Utah arrays and HF-LFPs was apt for characterizing motor behavior, we repeated our motor decoding analyses after artificially reducing the resolution of the HF-LFP signal via PCA. We found that taking linear projections of intra-region HF-LFPs yielded a significant decrease in classification accuracy, suggesting that intra-region population-level activity on small spatial scales is complex and possibly non-linear. Altogether, these findings show that HF-LFPs effectively captures significant aspects of intra-region population dynamics underlying reach-to-grasp motor states.

### How informative is intra-region heterogeneity?

We’ve shown that intra-region heterogeneity varies with neuromotor state and brain region. The underlying reasons for this, however, require further exploration. Firstly, since the EV, EC, and ΔPC metrics all involve a comparison to Baseline, their values could merely reflect involvement of the brain area during a particular neuromotor state. We believe, however, that much more can be gleaned from these metrics regarding the degree and type of region-level recruitment.

Latent dimensionality, for example, has been shown to be an informative measure of neural activity. Complex and naturalistic behaviors have been associated with a higher latent dimensionality (Altan, 2023; Mitchell 2023). Moreover, neuromotor states that imply finer motor control (e.g., reaching, grasping) or more external inputs may require coordination between a larger number of neural ensembles, resulting in a higher latent dimensionality. In the case of single unit activity, latent dimensionality represents the number of distinct patterns of coordinated neural activity within a population (i.e., neural ensembles; Gallego 2017). Analogously, since HF-LFP time series capture activity from a small number of populations, as opposed to the spiking of a single neuron, dimensionality in the case of HF-LFPs is likely proportional to the number of coordinated ensembles of neural populations underlying a neural state (Gallego-Carracedo 2024). Therefore, ΔPC should be proportional to a change in the number of ensembles of neural populations exhibiting modulated activity with respect to Baseline.

In addition to ΔPC, EV and EC served as complementary metrics of intra-region heterogeneity. Low EC, or low directional bias, indicates a balanced mixture of positive and negative neural responses relative to Baseline. This suggests a fine-grained spatial organization and diverse local micro-circuitry (Muthuswamy 2005; Gaunt 2011). By contrast, high EC implies a more region-wide, uniform response.

While EC is a measure of directional agreement, EV is a heterogeneity-of-magnitude metric (see Fig 1). High EV indicates a large spread of HF-LFP amplitudes across electrodes and, thus, reflects strong spatial selectivity or “hotspots” of engagement within a region. This metric is particularly informative in combination with EC (see Table 3). Low EV and EC indicate marginal involvement of an area (i.e., activity resembles Baseline). Low EV and high EC suggest a region-wide, coherent modulation consistent with states involving broad activation of the map (e.g., arousal). Monkey 1 cM1 exemplifies this (Fig 4). High EV and low EC imply participation of the region in multiple competing or complementary computations, consistent with higher order cognitive processes. This is consistent with cPMd in Monkey 1 (Fig 4). Finally, the combination of high EV and high EC reflects a coherent directional bias, either inhibitory or excitatory, along with local hotspots of activity. The best example of this in our findings is M1 in Monkey 2 which exhibited high EC and high EV for the Reach and Grasp states, respectively. Nonetheless, a larger cohort will be required to establish reliable group-level effects, since our metrics differed markedly between subjects.

**Table 3.** Heterogeneity metrics and their interpretation.

| EV | EC | Interpretation |
|---|---|---|
| Low | Low | Weak engagement of region in current behavior |
| Low | High | Region-wide, coherent modulation |
| High | Low | Balanced combination of strongly driven and inhibited populations |
| High | High | Strong directional bias with strong local specialization |

It is worth noting that we analyzed activity from cortical areas known to contain functional subregions due to the uneven distribution of their numerous cortical and thalamic inputs (Dea 2016; Strick 2021; Genon 2017; Genon 2018; He 1993). Future work will have to determine the relationship between the number of functional subregions and the heterogeneity metrics described in the present work. PMv, for instance, did not exhibit a high heterogeneity despite being known to contain at least five functional sub-regions (Kurata 2018). Moreover, it consistently ranked lower than the PMd and M1 in terms of neuromotor decoding performance. A better explanation of the informativeness of intra-region heterogeneity might be differences in the fine-grained spatial organization of the hand, digit, and arm representations between regions.

### Mesoscale heterogeneity and LFPs

Mesoscale heterogeneity of neural activity was previously investigated in resting state fMRI (rfMRI) studies. One specific metric, regional homogeneity (ReHo), has been found to covary with cognitive control, intelligence, and neurodevelopmental disorders (Jiang and Zuo 2016, Shen 2024). The metric, introduced in 2004 by Zang et al, was computed by taking Kendall’s coefficient of concordance between a given voxel and its neighbors (Zang 2004). Proponents state that this metric provides a novel perspective on local functional homogeneity and that it is an inherently multimodal and multiscale neuroimaging marker. Moreover, it has been shown to serve as a metric of network centrality (Jiang 2016; Shen 2024). Though ReHo has proven to be an effective metric, we have shown that EV, EC, and ΔPC also capture informative aspects of mesoscale activity. ReHo is typically computed directly from the time series of BOLD activity. EV and EC, however, can easily be applied to any features extracted from neurophysiological time-series. Moreover, EV and EC characterize heterogeneity by providing a metric of the spread and coherence of the data sources (i.e. channels, voxels, etc.) associated with a particular neural state, relative to another. The present work also differs from the aforementioned in that LFPs were used in lieu of rfMRI. The high temporal resolution of LFP enables investigation of rapidly evolving neuro-motor states, as found in a reach-to-grasp movement. Moreover, though some of the previous work involved the mesoscale, some did not. Ultra-high field strength, often in excess of 7 Tesla, is required to reach such spatial resolutions. In the present work, data was derived from Utah micro-electrode arrays with an inter-electrode spacing of 0.4mm and analyses were centered around the high frequency bands, the least correlated component of the signal. To our knowledge, this is the first time intra-region heterogeneity has been quantitatively investigated with LFPs.

### Heterogeneity is multifaceted

The present work demonstrates the multifaceted nature of intra-region heterogeneity. Although the trend is for more heterogeneous regions to yield better classification performance (better coding), this was not always the case. Monkey 1 cM1, for example, yielded high classification accuracy despite its higher than average intra-region correlation and average heterogeneity, according to our three metrics. cPMd in Monkey 1, in contrast, was the most heterogeneous region and boasted the best classification performance of any region. What is more, heterogeneity was informative even in relatively homogeneous regions. Heterogeneity of cM1 in Monkey 1 yielded the best classification performance of any region despite its high homogeneity (Fig 6). Neither intra-region correlation nor any of the three metrics used in this paper provide a full characterization of each region’s heterogeneity. Each metric captures a distinct aspect of heterogeneity, and further research will be required to determine which metrics capture the most informative aspects of intra-region neural dynamics.

### Population-level as a mesoscopic descriptive ground

We have shown that HF-LFPs capture complex population-level dynamics on small spatiotemporal scales (Fig 3), and that neuromotor states are encoded in the intra-region heterogeneity of neural population activity (Fig 6). Moreover, LFPs are known to be a more stable signal than spiking activity, and have been used to successfully reproduce several results previously based on SUA (Sharma 2015; Trautmann 2019). The combination of these findings support HF-LFP as a suitable candidate for multi-region analysis in systems neuroscience. With recent advances in electrophysiology such as the neuro-pixel technology, it is now possible to record from a large number of neural populations across brain regions and layers. Though certain contexts necessitate the use of single-unit spiking activity, population-level analyses may serve as an appropriate middle ground between SUA and broader signals, to characterize systems of coordinated neural populations in a functionally informative fashion.

### Potential Implications for BCI

Previous studies showed that motor states could be decoded from low to middle frequency LFPs (<250Hz) in BCI contexts (Heldman 2006). The present work extends and reinforces those results by demonstrating high performance decoding of sequential motor states from HF-LFP (200-500Hz) is possible even with short sample duration (0.25s samples) and few trials (40 & 25 trials for Monkeys 1 & 2, respectively), despite the 1/f power decay law. Specifically, our results show that resting, preparation, and execution states could be decoded from cPMd activity in a robust and reliable fashion from LFPs. This is particularly relevant for multi-array systems, since our findings suggest the cPMd as the best candidate for “high level” state decoding. This could then be used to select “low level” decoders trained on activity from other regions for finer control (e.g., fine finger coordination). This type of “gated” motor classification framework has already been shown to increase decoding performance and to reduce unintended movement (Aggarwal 2013, Xiao 2026). Our finding that intra-region heterogeneity is predictive of decoding performance could also inform array location selection. 7-Tesla fMRI, for example, can capture neural activity on a spatial scale comparable to Utah arrays and recent work has already identified it as an effective guide for pre-implantation mapping of task-related activity (Kromm 2026). Although future research will have to be performed to corroborate this, in principle, EV, EC, and ΔPC can be calculated from BOLD activity and be used to help inform array location.

## Conflict of interest statement

The authors declare no competing financial interests.

## Acknowledgments

This work was supported by the “la Caixa” Foundation’s doctoral INPhINIT fellowship for MD (LCF/BQ/DR22/11950018), and by project PID2023-149210NB-I00 by the Ministry of Science, Innovation and Universities of Spain for IC.

## Footnotes

1 Note we report the test statistic and p-value upper bounds obtained from multiple tests

## Notes

### Competing Interest Statement

The authors have declared no competing interest.

