## Supplemental Tables and Figures for "An LFP-based characterization of population-level heterogeneity during motor preparation & execution"

#### Tables

| | | | # of Electrodes | |
|---|---|---|---|---|
| Animal | Region | Array | p_max=0.05 | p_max=0.005 |
| Monkey 1 | cPMv | 64 | 61 | 45 |
| | cPMd | 32 | 32 | 30 |
| | iPMd | 64 | 61 | 44 |
| | iPMv | 64 | 64 | 64 |
| | cM1 | 32 | 32 | 32 |
| Monkey 2 | cPMv | 64 | 59 | 48 |
| | iPMd | 64 | 64 | 60 |
| | iPMv | 64 | 60 | 55 |
| | cM1 | 64 | 63 | 60 |

Table S1. Number of electrodes included in analyses. P-value columns indicate the number of electrodes that yielded at least one p-value below 0.05 or 0.005 when comparing mean classification accuracy to chance-level (see Methods).

| | *Movement Time (Monkey 1)* | | | | | *Movement Time (Monkey 2)* | | | |
|---|---|---|---|---|---|---|---|---|---|
| | cPMv | cPMd | iPMd | iPMv | cM1 | cPMv | iPMd | iPMv | cM1 |
| SA1 | 0.32 | 0.19 | -0.98$^{***}$ | 0.18 | -0.24 | -0.25 | 0.19 | 0.04 | -0.67 |
| | (0.20) | (0.33) | (0.36) | (0.22) | (0.24) | (0.21) | (0.17) | (0.03) | (0.39) |
| SA2 | -0.64$^{**}$ | 0.33 | 0.78$^{**}$ | -0.31 | | 0.05$^{***}$ | -0.55$^{**}$ | | -0.39 |
| | (0.31) | (0.30) | (0.36) | (0.24) | | (0.02) | (0.23) | | (0.23) |
| SA3 | 0.32 | 0.46 | -0.39 | -0.27 | | | | | |
| | (0.19) | (0.29) | (0.37) | (0.23) | | | | | |
| SA4 | | -0.30 | 0.59$^{**}$ | 0.19 | | | | | |
| | | (0.29) | (0.29) | (0.18) | | | | | |
| SA5 | | 0.85$^{**}$ | | 0.26 | | | | | |
| | | (0.41) | | (0.21) | | | | | |
| SA6 | | -0.63$^{**}$ | | -0.35 | | | | | |
| | | (0.27) | | (0.22) | | | | | |
| SA7 | | -0.73$^{**}$ | | | | | | | |
| | | (0.33) | | | | | | | |
| SA8 | | 0.57 | | | | | | | |
| | | (0.37) | | | | | | | |
| SA9 | | 0.51 | | | | | | | |
| | | (0.35) | | | | | | | |
| const | 0.54$^{*}$ | -0.72 | 0.57 | 0.84$^{***}$ | 0.79$^{***}$ | 0.70$^{***}$ | 0.85$^{***}$ | 0.46$^{***}$ | 1.58$^{***}$ |
| | (0.30) | (0.69) | (0.56) | (0.30) | (0.24) | (0.22) | (0.29) | (0.05) | (0.47) |
| Observations | 40 | 40 | 40 | 40 | 40 | 25 | 25 | 25 | 25 |
| $R^2$ | 0.17 | 0.51 | 0.28 | 0.24 | 0.02 | 0.31 | 0.26 | 0.08 | 0.20 |
| Adjusted $R^2$ | 0.10 | 0.36 | 0.19 | 0.10 | -0.00 | 0.25 | 0.19 | 0.04 | 0.13 |
| Residual Std. Error | 0.15 | 0.12 | 0.14 | 0.15 | 0.15 | 0.13 | 0.13 | 0.14 | 0.14 |
| F Statistic | 2.48$^{*}$ | 3.48$^{***}$ | 3.34$^{**}$ | 1.73 | 0.93 | 5.05$^{**}$ | 3.81$^{**}$ | 2.09 | 2.75$^{*}$ |
| Note: | | | | | | $^{*}$p<0.1; $^{**}$p<0.05; $^{***}$p<0.01 | | | |

Table S2. Movement time vs. HF-LFP-derived spectral amplitude (SA) regression results during Pre-Reach. A linear regression model was fit to spectral amplitude (SA) from each region during the Pre-Reach period (e.g., 5 models were it to each of the 5 regions recorded from Monkey 1). The number of variables (i.e., electrodes) to include in each model was determined via forward stepwise regression. The subset of electrodes included in each model was the one yielding the highest adjusted $R^2$.

| | *Movement Time (Monkey 1)* | | | | | *Movement Time (Monkey 2)* | | | |
|---|---|---|---|---|---|---|---|---|---|
| | cPMv | cPMd | iPMd | iPMv | cM1 | cPMv | iPMd | iPMv | cM1 |
| SA1 | 0.38 | $-0.92^{**}$ | 0.42 | 0.20 | -0.22 | -0.02 | $-0.72^{**}$ | $-0.99^{**}$ | -0.23 |
| | (0.23) | (0.34) | (0.33) | (0.17) | (0.20) | (0.01) | (0.32) | (0.40) | (0.22) |
| SA2 | | 0.54 | 0.60 | | -0.31 | $0.05^{**}$ | -0.28 | | $-0.32^{**}$ |
| | | (0.36) | (0.42) | | (0.18) | (0.02) | (0.27) | | (0.14) |
| SA3 | | 0.16 | 0.43 | | | | $1.04^{**}$ | | $-0.47^{***}$ |
| | | (0.34) | (0.37) | | | | (0.43) | | (0.14) |
| SA4 | | $-0.51^{**}$ | | | | | | | |
| | | (0.25) | | | | | | | |
| SA5 | | $-0.56^{*}$ | | | | | | | |
| | | (0.33) | | | | | | | |
| SA6 | | $0.51^{*}$ | | | | | | | |
| | | (0.30) | | | | | | | |
| SA7 | | $0.51^{*}$ | | | | | | | |
| | | (0.25) | | | | | | | |
| const | 0.16 | 0.84 | $-0.90^{*}$ | $0.35^{*}$ | $1.07^{***}$ | $0.49^{***}$ | 0.47 | $1.50^{***}$ | $1.55^{***}$ |
| | (0.24) | (0.64) | (0.51) | (0.18) | (0.24) | (0.04) | (0.52) | (0.40) | (0.28) |
| Observations | 40 | 40 | 40 | 40 | 40 | 25 | 25 | 25 | 25 |
| $R^2$ | 0.06 | 0.39 | 0.22 | 0.03 | 0.12 | 0.20 | 0.30 | 0.21 | 0.53 |
| Adjusted $R^2$ | 0.04 | 0.26 | 0.15 | 0.01 | 0.07 | 0.13 | 0.20 | 0.18 | 0.46 |
| Residual Std. Error | 0.15 | 0.13 | 0.14 | 0.15 | 0.15 | 0.14 | 0.13 | 0.13 | 0.11 |
| F Statistic | 2.62 | $2.98^{**}$ | $3.31^{**}$ | 1.31 | $2.51^{*}$ | $2.72^{*}$ | $2.95^{*}$ | $6.11^{**}$ | $7.83^{***}$ |
| Note: | | | | | | $^{*}p<0.1$; $^{**}p<0.05$; $^{***}p<0.01$ | | | |

Table S3. Movement time vs. HF-LFP-derived spectral amplitude (SA) regression results during Reach. A linear regression model was fit to spectral amplitude (SA) from each region during the Reach period (e.g., 5 models were it to each of the 5 regions recorded from Monkey 1). The number of variables (i.e., electrodes) to include in each model was determined via forward stepwise regression. The subset of electrodes included in each model was the one yielding the highest adjusted $R^2$.

| | | $\Delta PC$ | | | |
|---|---|---|---|---|---|
| | | Explained Variance | | | |
| Region | State | 0.60 | 0.66 | 0.70 | 0.75 |
| cPMv | Pre-Reach | 0 | 0 | 0 | 0 |
| | Reach | 0.5 | 0 | 0.5 | 1 |
| | Grasp | 0 | 0 | 1 | 1 |
| | Post-Grasp | 1 | 1 | 1 | 1 |
| cPMd | Pre-Reach | 0 | 0 | 0 | 0 |
| | Reach | 1 | 1 | 1 | 0.5 |
| | Grasp | 1 | 1 | 1 | 1 |
| | Post-Grasp | 1 | 1 | 1 | 1 |
| iPMd | Pre-Reach | 0 | 0 | 0.5 | 1 |
| | Reach | 1 | 2 | 2 | 2 |
| | Grasp | 1 | 1.5 | 1.5 | 1.5 |
| | Post-Grasp | 1 | 2 | 2 | 2.5 |
| iPMv | Pre-Reach | 0 | 0 | 0 | 0 |
| | Reach | 0 | 0 | 0 | 0 |
| | Grasp | 0 | 0 | 0 | 0 |
| | Post-Grasp | 0 | 0 | 0 | 0 |
| cM1 | Pre-Reach | 0 | 1 | 1 | 0.5 |
| | Reach | 2 | 1.5 | 1 | 1.5 |
| | Grasp | 1 | 1 | 1 | 1 |
| | Post-Grasp | 1 | 1 | 1 | 1 |

Table S4. ΔPC stability across explained variance thresholds for Monkey 1. ΔPC values reflect the median (across trials) difference in the number of principal components required to explain the corresponding explained variance threshold. All analyses used the 0.66 threshold.

| | *Sensitivity* | |
|---|---|---|
| | Monkey 1 | Monkey 2 |
| Group Var | 4.25 | 0.00 |
| | (3.86) | (0.33) |
| Intercept | 0.41 | $0.40^{*}$ |
| | (0.22) | (0.17) |
| EC | -0.04 | 0.12 |
| | (0.23) | (0.21) |
| EV | 0.16 | $0.28^{*}$ |
| | (0.16) | (0.11) |
| ΔPC | $0.12^{**}$ | $0.13^{**}$ |
| | (0.04) | (0.05) |
| Observations | 20 | 16 |
| Residual Std. Error | 0.09 | 0.15 |
| Note: | $^{*}$p<0.05; $^{**}$p<0.01; $^{***}$p<0.001 | |

Table S5. Multiple linear regression coefficients from models predicting sensitivity from three metrics of intra-region heterogeneity: Effect Coherence (EC), Effect Variance (EV), and latent dimensionality (ΔPC). The single-feature classification accuracy p-value cutoff (see Methods) for an electrode to be included in the analysis was reduced from 0.05 to 0.005. Sensitivity is a measure of classification (i.e. decoding) performance which indicates the proportion of true positives to actual positives.

### Figures

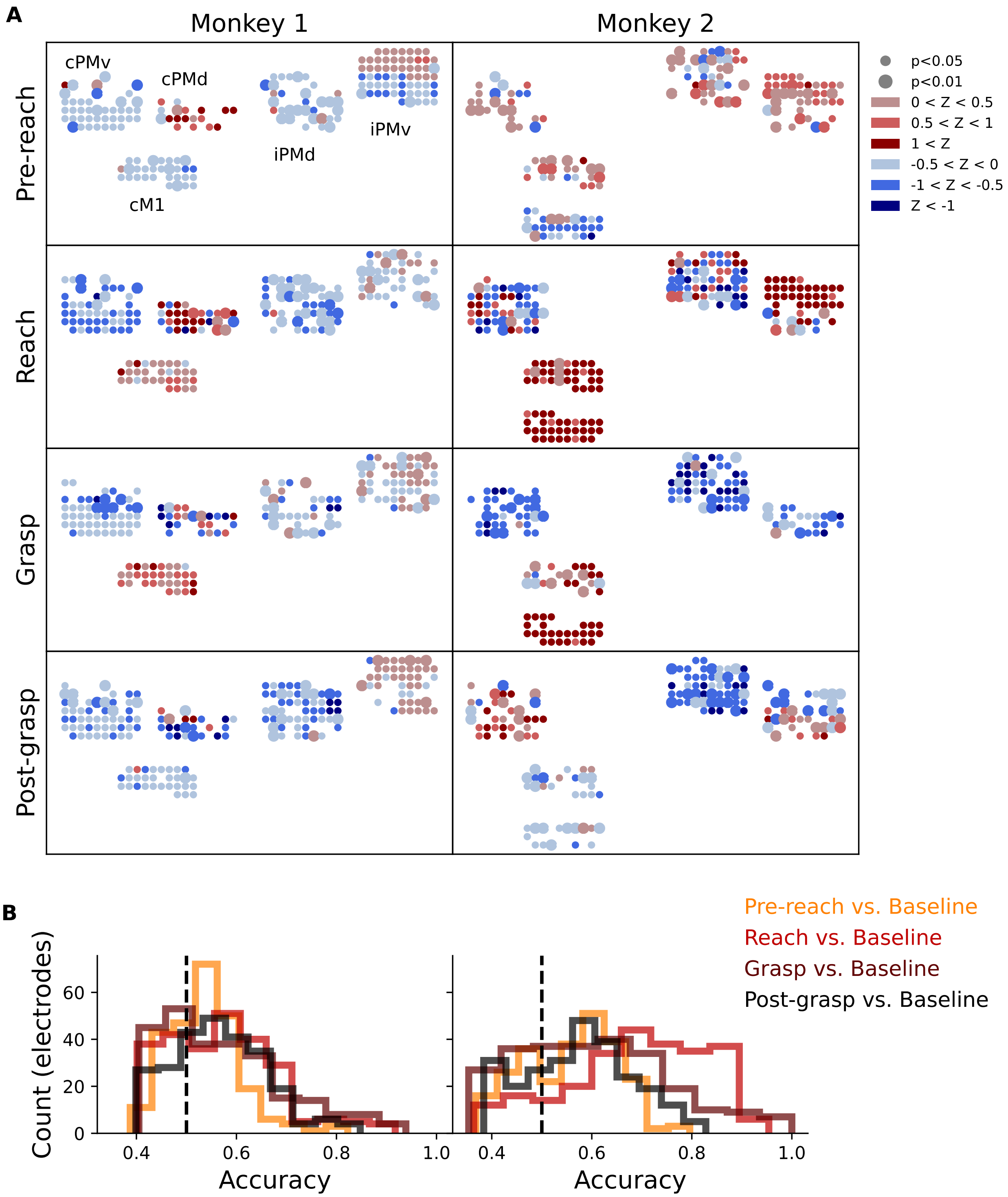


Figure S1. Single-feature binary classification results. Binary classification was performed between Baseline and each post-Baseline state using the spectral amplitude features derived from the 200-500 Hz band. **A)** Classification results scatter plots. Marker color indicates the modified Z-score relative to Baseline. Marker radius indicates p-value of the binary classification accuracy compared to chance-level (50%). Electrodes which did not exhibit above-chance classification accuracy are not shown. **B)** Classification accuracy histograms. The dashed vertical line represents chance-level (50%).

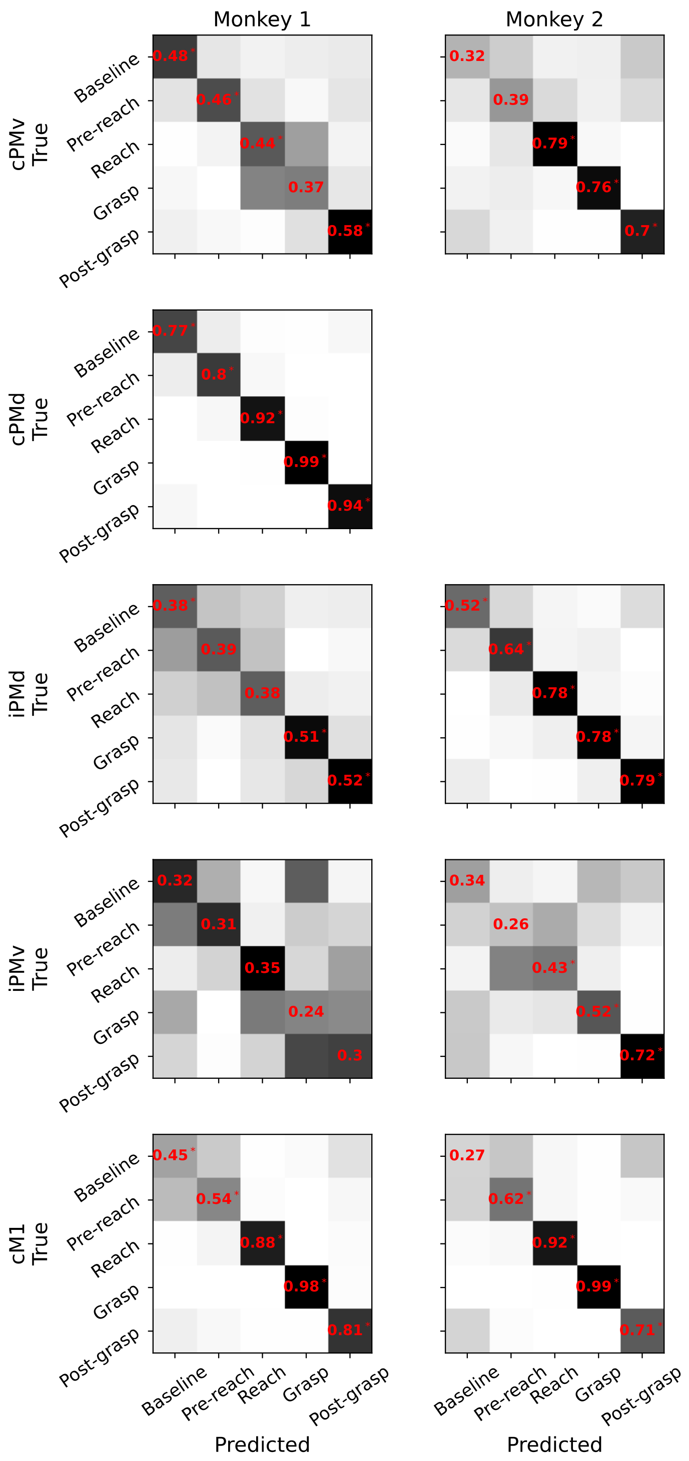


Figure S2. By-region classification results heatmaps. By-region, row-normalized confusion matrices for Monkey 1 (left, and Monkey 2 (right). By-class sensitivity is shown in red on the diagonal and statistical markers indicate significance as determined by permutation testing. Significance is only reported on the diagonal.

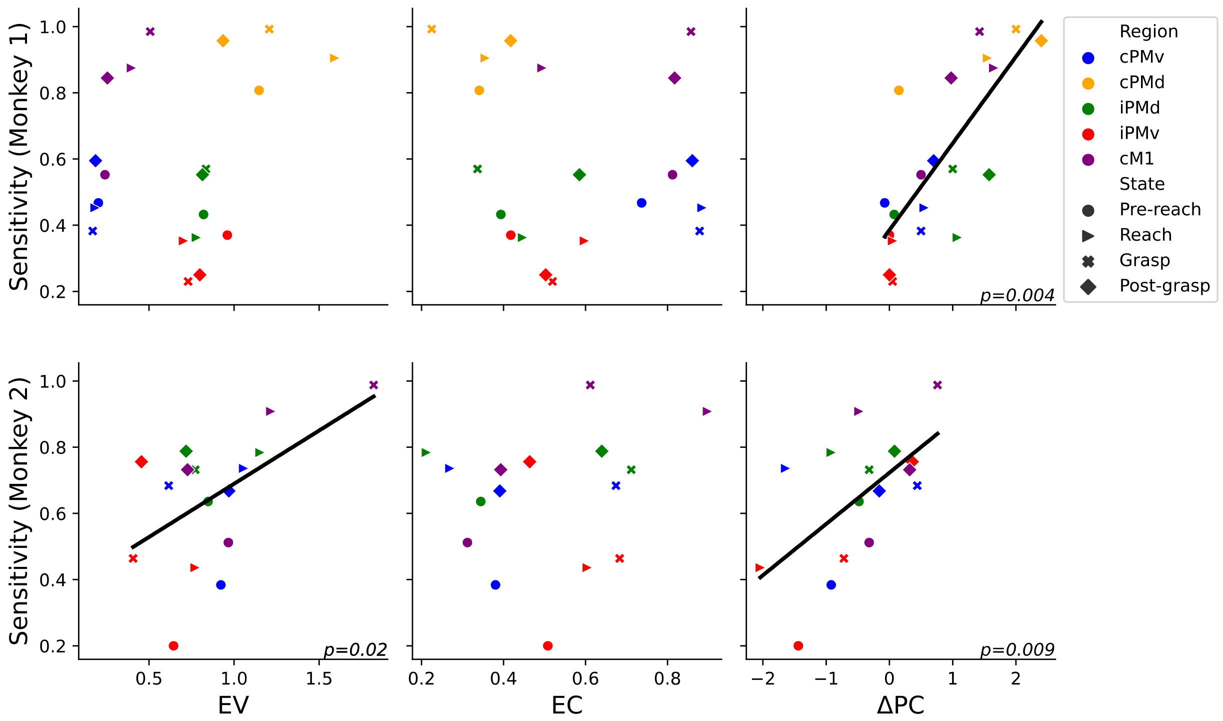


Figure S3. Sensitivity as a function of intra-region heterogeneity (EV, EC, & ΔPC) of HF-LFP for Monkey 1 (top row) and Monkey 2 (bottom row). The single-feature classification accuracy p-value cutoff (see Methods) for an electrode to be included in the analysis was reduced from 0.05 to 0.005. Colors represent brain areas and shapes represent neuromotor states. Chance-level is 0.2.
